# Oligodendroglial deletion of the microcephaly gene Cit-k disrupts cortical connectivity and cognitive function

**DOI:** 10.64898/2026.08.07.743469

**Authors:** Martino Bonato, Federica Marchiotto, Maryam Khastkhodaei Ardakani, Federico G. P. Ferrari, Niccolò Di Cintio, Annamaria Renna, Ottavia Maria Roggero, Francesca Montarolo, Valentina Cerrato, Angelisa Frasca, Benedetto Sacchetti, Annalisa Buffo, Marco Cambiaghi, Enrica Boda

## Abstract

Neurodevelopmental disorders (NDDs) are increasingly recognized as disorders of brain connectivity and circuit dysfunction. Growing evidence suggests that glial cell and myelin abnormalities may actively contribute to these alterations. Yet, they have been often considered secondary consequences of impaired neuronal development rather than primary drivers of circuit dysfunction. Primary autosomal recessive microcephaly type 17 (MCPH17) is a severe NDD caused by mutations in the *CIT* gene, encoding Citron kinase (CIT-K). The disease is associated with cognitive and motor deficits, epilepsy susceptibility, and marked hypomyelination in both patients and mouse models, suggesting a contribution of oligodendroglial dysfunction to disease pathophysiology.

Here, we investigated the specific role of oligodendroglial Cit-k loss using Sox10^Cre^;Cit-k^fl/fl^ mice, in which Cit-k is selectively deleted in oligodendrocyte-lineage cells. Mutant mice displayed impaired forebrain myelination at juvenile stages and persistent cortical hypomyelination in adulthood. Despite preserved gross motor function, adult mutants showed deficits in fine motor control, working and recognition memory, and auditory fear memory. These impairments were associated with altered cortico-cortical and cortico-hippocampal functional connectivity. Moreover, consistent with the clinical MCPH17 phenotype, mutant mice exhibited increased susceptibility to kainate-induced seizures.

Together, our findings show that oligodendroglial Cit-k loss and the resulting hypomyelination are sufficient to produce long-lasting neurological and behavioral impairments independently of primary neuronal defects. These results identify oligodendrocytes as active contributors to MCPH17 and support a broader role for myelin abnormalities in NDDs.

**Highlights:**

- Cit-k deletion in oligodendroglia disrupts forebrain myelination
- Cortical hypomyelination persists in adult mutant mice
- Mutant mice show deficits in motor control and memory
- Cortico-cortical and cortico-hippocampal connectivity are altered
- ligodendrocytes contribute to microcephaly-associated dysfunctions

## Introduction

Neurodevelopmental disorders (NDDs) comprise a heterogeneous group of congenital or early-acquired conditions that disrupt the formation and maturation of the central nervous system (CNS), leading to cognitive, motor, and behavioral impairments (Silbereis et al., 2016). Traditionally, the pathogenesis of these disorders has been interpreted through a neuron-centric perspective, with clinical phenotypes attributed to intrinsic neuronal defects, including impaired neural progenitor cell (NPC) proliferation, abnormal neuronal migration, defective differentiation, or altered synaptic connectivity (Rakic, 2009; Subramanian et al., 2020). This view has been particularly dominant in early NDDs such as microcephaly, which has been regarded as a group of prototypical “neuronal disorders” because of their association with impaired neurogenesis and early brain growth failure (Jayaraman et al., 2018). Accordingly, glial abnormalities - most commonly hypomyelination (Dupuis et al., 2015; Nakayama et al., 2015; Fujikura et al., 2013; Ogi et al., 2018; Tisoncik-Go et al., 2024) - have long been considered secondary consequences of primary neuronal defects rather than active contributors to disease pathogenesis.

Among NDDs, primary hereditary microcephaly (MCPH) best exemplifies this neuron-centric perspective. MCPH is mainly caused by germline mutations in genes required for early corticogenesis, including MCPH1, WDR62, ASPM, and CIT (Jayaraman et al., 2018; Gilmore and Walsh, 2013; Bianchi et al., 2020). These genes encode proteins involved in key processes regulating NPC proliferation and genome integrity, such as mitotic progression, centrosome dynamics, chromosome segregation, and DNA replication/repair. Their disruption impairs NPC survival and expansion during embryonic development, ultimately reducing the generation of both neuronal and glial progeny (Siskos et al., 2021). As a consequence, MCPH patients display variable degrees of brain volume reduction and cortical malformations, frequently associated with cognitive and motor deficits, epilepsy, sensory impairments, and, in severe cases, early lethality (Mahmood et al., 2011).

Among genes implicated in MCPH, CIT has been associated with particularly severe neurodevelopmental phenotypes. Biallelic mutations in CIT, encoding the serine/threonine kinase Citron kinase (CIT-K), cause primary autosomal recessive microcephaly type 17 (MCPH17), a rare disorder characterized by reduced brain size, simplified gyrification, ventriculomegaly and variable neurological impairments, including cognitive, language, and motor deficits, epileptic seizures and even neonatal lethality (Li et al., 2016; Harding et al., 2016; Shaheen et al., 2016; Basit et al., 2016; <u>Microcephaly 17, Primary, Autosomal Recessive - MalaCards</u>). Consistent with human pathology, *Cit-k* knockout (KO) mice display dramatic reductions in cortical, hippocampal, and cerebellar structures, associated with ataxia, seizures, and early mortality within the first postnatal weeks (Di Cunto et al., 2000). Mechanistically, CIT-K plays essential roles in cytokinesis, cytoskeletal organization, and DNA damage repair (Di Cunto et al., 2000; Bianchi et al., 2017; Iegiani et al., 2025). Accordingly, loss of Cit-k function results in defective NPC division, accumulation of DNA damage, premature cell-cycle exit, and apoptosis (Bianchi et al., 2020; Pallavicini et al., 2024; Iegiani et al., 2025), ultimately leading to reduced neuronal populations (Di Cunto et al., 2000; Muzzi et al., 2009). In addition to neuronal defects, our previous work revealed severe hypomyelination in both Cit-k KO mouse brain and cortical tissue from an MCPH17 patient, suggesting that oligodendroglial dysfunction may represent an important and previously overlooked component of disease pathogenesis (Boda et al., 2022). In Cit-k KO mice, CNS hypomyelination was not caused by impaired generation of oligodendrocyte lineage cells, but rather by increased apoptosis and senescence of oligodendrocyte progenitor cells (OPCs) accumulating DNA damage. Consistent with these findings, the cortex of an MCPH17 patient showed an approximately 50% reduction in OPC density. Importantly, oligodendroglia abnormalities – i.e. hypomyelination, OPC apoptosis and senescence - were also observed in juvenile Sox10^Cre^;Cit-k^fl/fl^ mice, in which Cit-k deletion was restricted to oligodendroglial cells (Boda et al., 2022). Together, these findings support a cell-autonomous role for CIT-K in oligodendrocyte lineage cells and suggest that myelination defects can develop independently of primary neuronal defects.

In the CNS, myelination is essential for efficient action potential conduction and for the development of proper motor, sensory, and cognitive functions (Stadelmann et al., 2019; Khelfaoui et al., 2024). Oligodendroglial cells also support neuronal function through metabolic coupling and regulation of neuronal excitability (Schirmer et al., 2018; Simons et al., 2024), being increasingly recognized as key regulators of circuit assembly and maturation, neuronal synchrony, and higher-order cognitive functions, including learning and memory (Khelfaoui et al., 2024; Xin and Chan, 2020). Consistent with this, developmental abnormalities in oligodendroglia and myelination have recently been implicated in multiple neurodevelopmental conditions, including autism spectrum disorders and intellectual disability syndromes, suggesting that myelin/oligodendroglia dysfunction may contribute to disease genesis, evolution and neurological outcome (Khelfaoui et al., 2024; Kimchi-Feldhorn et al., 2026; Ramesh et al., 2025).

To define the specific contribution of oligodendroglial abnormalities to the pathogenesis of MCPH17, here we took advantage of the conditional Sox10^Cre^;Cit-k^fl/fl^mouse model and investigated how oligodendrocyte-specific loss of Cit-k affects CNS myelination, behavioral outcomes and cortical connectivity at juvenile and adult stages. Results show that oligodendroglia-specific loss of Cit-k cell-autonomously disrupts myelin deposition throughout the juvenile forebrain. Although myelination partially recovers over time, pronounced hypomyelination persists predominantly in the adult cerebral cortex, revealing a selective and long-lasting vulnerability of cortical regions. These structural abnormalities are accompanied by functional alterations. In adulthood, mutant mice exhibit impairments in movement initiation and accuracy, as well as deficits in working memory, short-term memory, and associative fear learning, despite the partial recovery of myelination. These behavioral alterations are associated with disrupted cortico-cortical and cortico-hippocampal functional connectivity during task execution. Importantly, mutant mice were also more susceptible to kainate-induced seizures, consistent with seizure occurrence in a subset of MCPH17 patients.

Collectively, these findings indicate that oligodendroglial Cit-k loss and the resulting hypomyelination are sufficient to drive persistent cognitive and network dysfunctions, supporting an active contribution of myelin/oligodendroglial abnormalities to the pathophysiology and long-term neurological manifestations of MCPH17.

## Material and Methods

### Experimental animals

The conditional Sox10^Cre^;Cit-k^fl/fl^ mouse line was used to selectively delete Cit-k in oligodendroglia (Boda et al., 2022). To obtain this mouse line, Cit-k^fl/fl^ mice (Pallavicini et al., 2018), originally from UC Davis KOMP repository as Cittm1a(KOMP)Wtsi, were crossed with Sox10^Cre^ (Matsuoka et al., 2005; i.e. B6;CBA-Tg(Sox10-cre)1Wdr/J, The Jackson Laboratory). Age-matched Cre-carrying (i.e. Sox10^Cre^ and Sox10^Cre^;Cit-k^fl/+^) mouse littermates have been used as controls (Ctrl). Mice were housed in the vivarium under standard conditions (12-hr light/12-hr dark cycle at 21°C) with food and water ad libitum. The project was designed according to the guidelines of the NIH, the European Communities Council (2010/63/EU) and the Italian Law for Care and Use of Experimental Animals (DL26/2014). It was also approved by the Italian Ministry of Health (authorization #589/2023-PR to AB). The study was conducted according to the ARRIVE guidelines.

Histological and behavioral analyses have been performed in juvenile (i.e. postnatal day 14, P14) and adult (3-5 months) mice. Isolation-induced ultrasonic vocalizations (USVs) were recorded exclusively at P10 and P14. Extracellular local field potential (LFP) analyses have been performed exclusively in 3-5 months-old mice. Since MCPH17 affects both males and females and, due to its rarity, possible sex-dependent differences in disease manifestations remain unclear, except for fear conditioning experiments (see below), both female and male mice have been included in the experiments.

For histological analyses, animals were deeply anesthetized (ketamine-xylazine 90 mg/kg and 9 mg/kg, respectively, i.p.) and transcardially perfused with 4% paraformaldehyde (PFA; Sigma-Aldrich) in 0.1 M phosphate buffer (PB). For western blotting, animals were deeply anesthetized (as above), and brains were quickly removed and manually dissected. All samples were rapidly frozen in −80°C liquid nitrogen and then stored at −80°C.

### Histological procedures

After mouse perfusion, brains were postfixed overnight, cryoprotected, and processed according to standard immunohistological procedures (Boda et al., 2022). Brains were cut in 30 μm thick coronal sections collected in PBS and then stained to detect the expression of different antigens: NG2 (rb, 1:200, Sigma-Aldrich AB5320); BCAS1 (rb, 1:500, Synaptic Systems); CC1 (m, 1:100, Anti-APC [CC-1], Abcam); MBP (m, Smi-99 clone, 1:1000, Biolegend). Incubation with primary antibodies was made overnight at 4 °C in PBS with 1-2% Triton-X 100. The sections were then exposed for 3 h at room temperature (RT) to secondary Cy3, Cy5 (Jackson ImmunoResearch Laboratories) or Alexa Fluor 488, Alexa Fluor 555 (Molecular Probes) -conjugated antibodies. 4,6-diamidino-2-phenylindole (DAPI, Fluka) was used to counterstain cell nuclei. After processing, sections were mounted on microscope slides with Tris-glycerol supplemented with 10% Mowiol (Calbiochem). Myelin silver nitrate Gallyas staining was performed as in (Pistorio et al., 2006). Nissl staining was performed as in (Boda et al., 2022).

### Image acquisition and data analysis

For the neuroanatomical analyses, Nissl-stained slices were imaged with an AxioScan instrument (Carl Zeiss Microscopy, New York, United States) at 4x magnification. Images were processed using Zen 3.0 (Zen light) blue edition (Carl Zeiss Microscopy, New York, USA). Parameters including forebrain area (hemi-section), ratio of lateral ventricle area over forebrain area, thickness of motor (M1) cortex and corpus callosum (CC, sampled at the dorsal midline) were measured on images of the same rostro caudal level for Sox10^Cre^;Cit-k^fl/fl^*vs* Ctrl mice. The rostro-caudal distribution of myelination was assessed on serial coronal sections of the forebrain and on sagittal cerebellar and horizontal spinal cord slices, by whole-slice imaging of either MBP immunofluorescence or Gallyas-stained sections acquired at 10× magnification. Whole-section images were generated using the automatic stitching function of Neurolucida, which combines sequential overlapping image tiles into a single high-resolution reconstruction. Immunolabeled histological specimens were examined using a Nikon C1 confocal microscope (with the associated EZ-C1 Ver3.90 software, Nikon), or a Leica TCS SP5 confocal microscope (with the associated LAS AF 4.0 software, Leica Microsystems). Confocal images (1024 × 1024 pixels) were acquired at 20× and 40×. Adobe Photoshop 6.0 (Adobe Systems, San Jose, CA) was used to assemble the final plates. Quantitative evaluations were performed on confocal images with Fiji/ImageJ (Research Service Branch, National Institutes of Health, Bethesda, MD; available at http://rsb.info.nih.gov/ij/). To analyze the expression level of MBP, the positive fractioned area (i.e. the percentage of positive pixels) was quantified in 40× confocal image stacks comprising 20 optical slices 0.99 µm thick (for MBP). Myelination was also quantified on Gallyas-stained sections using whole-slice images acquired at 10× magnification. Regions of interest (ROIs) corresponding to the relevant brain areas were delineated in ImageJ, and the fraction of stained area within each ROI was measured after intensity thresholding. Density of oligodendroglial (NG2+, BCAS1+, CC1+) cells was calculated as number of cells per mm^2^ in 40× confocal image stacks comprising 20 optical slices 0.99 µm thick. At least three animals and at least three sections per animal have been analyzed for each experimental group.

### Western Blotting

Brain tissues were lysed in ice-cold RIPA buffer (300 mM NaCl, 100 mM Tris-HCl, 2% NP-40, 1% sodium deoxycholate, 10 mM EDTA, 0.2% SDS, pH 8) using a sonicator (30 kHz, 15 pulses). Homogenates were centrifuged for 30 minutes at 12,000 g at 4°C, and supernatants were collected. Protein quantification was measured by BCA assay (ThermoFisher Scientific; #23225). Samples containing 10 µg of proteins were prepared in sample buffer and heated at 95°C for 5 minutes. Samples were separated by SDS-PAGE on TGX stain-free gels (10% of acrylamide gradient, Bio-Rad, #1610183) and proteins were blotted onto a nitrocellulose membrane using a semidry transfer apparatus (Trans-Blot Semi-Dry Transfer Cell; Bio-Rad). Membranes were incubated 5 minutes in EveryBlot Blocking Buffer (Bio-Rad; #12010020) and then incubated overnight at 4°C with the following primary antibodies: anti-Smi312 (rb, 1:500 in EveryBlot Blocking Buffer; anti-Neurofilament pan axonal marker, clone #Smi312, Biolegend), anti-CNPase (m, 1:500 in EveryBlot Blocking Buffer; clone #11-5B, Abcam), anti-MBP (m, 1:1000 in EveryBlot Blocking Buffer; clone #Smi-99, Biolegend). After three washes in TBST (Tris- 5% nonfat milk and 0.1% Tween-20, pH 7.4), membranes were incubated with HRP-conjugated secondary antibody for 1 hours at RT (1:5,000; Jackson ImmunoResearch). The immunocomplexes were visualized by using the ECL substrate (Cyanagen) and Essential V6 imaging platform, UVITEC system (Cleaver Scientific Ltd). Band density measurements were performed using UVITEC software. Results were normalized to total protein content visualized by a TGX stain-free method (Bio-Rad).

### Behavioral characterization at juvenile stage

A battery of tests was employed to investigate mouse behavioral phenotype at juvenile stage, i.e. P14 (Feather Schussler & Ferguson, 2016). Body weight was measured before the tests, using a standard small animal balance (Suppl. Fig. 2A). The following parameters were taken into consideration:

1. *Motility*: In a 20 x 26 cm open field arena, mice were allowed to walk freely for 2-3 minutes while videorecorded by a camera placed on the ceiling. Using the open-source Behavioral Observation Research Interactive Software (BORIS, www.boris.unito.it, Friard & Gamba, 2016) software, the percentage of time the animals were in motion compared to the overall time was measured on the recorded videos.
2. *Maturity of the gait pattern (ambulation score)*: during the above-mentioned test, two independent operators scored mouse walking pattern on a scale of 0 to 2 (0 = crawling; 1 = transition; 1.5 = the mouse lifts its tail or walks only on the front part of the legs; 2 = the mouse raises its tail and walks only on the front part of the legs).
3. *Posture*: As a measure of a possible ataxic phenotype, we evaluated the hindlimb foot angle, as indicative of the acquisition of a compensatory posture, consisting of spreading the hindlimbs apart to maintain balance. This analysis was performed on the videos described above using the ImageJ software on 20 frames per mouse (with the apex on the tail when the animal is completely aligned).
4. *Negative geotaxis*: for the evaluation of motor coordination and vestibular system function, mice were placed on a 45° inclined smooth surface with the head facing down. The ability of the mouse to return to the anti-gravity position was evaluated within three consecutive tests (maximum time of 1 minute to carry out the test). The time required to complete the task was measured.
5. *Surface Righting:* Surface righting reflex was measured by placing juvenile mice on their back on a surface with their limbs close to the body. Mice were kept in position by an operator until they showed immobility for three seconds. Surface righting reflex time was measured with the start at the release of the limbs by the operator and end when the animal flips on its belly and all four paws touch the ground. Three consecutive trials were performed to obtain the mean reflex time for each animal.
6. *Hindlimb suspension test*: this test allowed to assess the strength of mouse hind limbs. Pups were placed head down, hanging by the hindlimbs into a 50 ml plastic tube padded with laboratory wipes for protection. The latency to fall from the edge of the tube was evaluated and a score was assigned to the hindlimb posture from 4 (normal hindlimb separation with tail raised) to 0 (constant clasping of the hindlimbs with the tail lowered). The test was performed in three consecutive trials and the average latency and score were calculated.
7. *Ultrasonic vocalizations (USVs) upon isolation*: isolation-induced USVs were recorded at P10 and P14 during the first hours of the light phase. Prior to testing, mice were maintained in the home cage and then individually isolated in a 15 × 10 cm recording chamber placed inside a sound-attenuating cubicle. USVs were recorded for 3 min using an Avisoft Bioacoustics UltraSoundGate CM16/CMPA microphone positioned 10 cm above the animal and connected to an UltraSoundGate 116Hb recording interface and Avisoft Recorder software. Audio files were analyzed using the MATLAB-based software DeepSqueak. Vocalizations were automatically detected and subsequently manually curated to exclude artifacts and ensure accurate contour identification and call classification. Acoustic features such as Contour Shape, Duration and Frequency were used as input in DeepSqueak to generate a UMAP embeddings of all recorded vocalizations, then exported for clustering and statistical analyses in RStudio.

### Behavioral characterization of adult mice

Behavioral assessment was performed in adult (3-5 months old) mice during the light phase by operators blind to genotype.

1. Visual function was assessed using the *visual placing reflex test*, as previously described for rodent sensorimotor evaluation (Schaar et al., 2010). Briefly, mice were gently moved toward a tabletop surface (either vertically or horizontally) and forelimb extension responses were evaluated as an index of visually guided motor behavior. Responses were scored according to the distance from the surface at which the animal initiated forepaw extension (0 - catatonic animal that did not respond even after direct contact with the surface; 1 - low response that occurs only when the animal comes in contact with the surface; 2 - slow response near the cage, within vibrissae contact distance; 3 - clear response from a medium distance (less than 2.5 cm); 4 - ready response at long distance (more than 2.5 cm)). Each mouse underwent three consecutive trials, and the average score was used for statistical analysis.
2. *Acoustic startle responses* were assessed in a sound-attenuated chamber using a custom AnyMaze-controlled protocol. Following 1 min habituation, mice were exposed to 11-kHz acoustic bursts of increasing intensity (70–100 dB, 5-dB increments, 30 s interstimulus interval). Behavioral responses were video-recorded and scored offline as positive or negative startle reactions. Startle sensitivity was expressed as the percentage of positive responses, as previously described for rodent acoustic startle assessment (Geyer et al., 2001).
3. Mouse *motility* in the open field arena was evaluated as described above for juvenile mice.
4. *Hindlimb clasping* behavior was assessed during tail suspension, as previously described for neurological phenotyping in mice (Guyenet et al., 2010). Mice were suspended by the tail for ∼10 s and video-recorded from a ventral view. Representative frames in which the animal was immobile and aligned with the camera were selected for analysis. Hindpaw distance was measured as an index of clasping behavior and normalized using fixed spatial references within the recording. Mean hindpaw distance values were calculated for each animal and normalized to control littermates.
5. *Motor initiation* and posture-release performance were assessed using a modified *bar test* adapted from previous studies in CNP-deficient mice (Hagemeyer et al., 2012). Briefly, mice were placed in a forced rearing posture with the forepaws positioned on a horizontal plastic bar (14 cm length, 0.3 cm diameter) suspended 5 cm above the cage floor, while the hindpaws remained on the ground. Trials were video-recorded and the latency to release the first and second forepaw from the bar was measured as an index of movement initiation ability. Each mouse underwent three consecutive trials and mean release latencies were used for statistical analysis.
6. *Motor coordination and learning* was evaluated using the accelerating *rotarod test* (Mouse Rota-Rod, Ugo Basile Biological Research Apparatus, Comerio, Italy), consisting of three daily sessions of three trials over three consecutive days (4–65 rpm over 300 s, inter-trial interval 60s). Latency to fall or passive rotation was recorded and averaged across trials (Hoxha et al., 2012).
7. *Gait* and *locomotion features* were assessed using the Noldus Catwalk system. Mice were tested in a dimly lit room, placed on the glass walkway and recorded until completion of 6 full compliant runs. CatWalk XT version 10.7 software was used for automated detection of footprints, manual curation of detections and motor parameters analysis.
8. *Fine motor coordination and balance* were assessed using the beam-walking test, as previously described (Hoxha et al., 2012). Mice were required to traverse a 1-m-long, 1-cm-wide elevated wooden beam leading to a dark enclosed escape box, while the starting platform was illuminated to provide an aversive stimulus. Animals were habituated to the apparatus before testing. Beam traversal time and hindlimb slips were recorded by two operators. Each mouse performed three trials per day over two consecutive days, and mean values were used for statistical analysis.
9. *Spatial working memory* was assessed using the spontaneous alternation *Y-maze test*, as previously described (Kraeuter et al., 2019). The apparatus consisted of three identical black plastic arms positioned at 120° angles from one another and surrounded by distal visual cues. Mice were placed individually in the maze and allowed to freely explore for 8 min while being video-recorded. Arm entry was scored when all four paws entered an arm. Spontaneous alternation performance was calculated as the percentage of consecutive entries into three different arms over the total number of alternation opportunities, and used as an index of spatial working memory.
10. *Short-term recognition memory* was assessed using the *novel object recognition (NOR)* test, as previously described (Bevins & Besheer, 2006). Mice were tested in a Noldus PhenoTyper arena using a three-phase protocol consisting of habituation, familiarization, and test sessions. During the familiarization phase, two identical objects were placed symmetrically within the arena, whereas in the test phase one familiar object was replaced with a novel object of similar dimensions but different shape and appearance. Mice were allowed to freely explore the arena for 15 min during each phase, with a 2.5 h intertrial interval. Object exploration was defined as direct contact and/or sniffing behavior. Recognition memory performance (i.e. preference for the novel object) was expressed as the percentage of exploration time directed toward the novel object over total object exploration time.
11. *Associative emotional memory* was assessed using an *auditory fear-conditioning* paradigm, as previously described (Cambiaghi et al., 2016, Concina et al., 2022). Adult male mice were used to minimize variability associated with the estrous cycle. During conditioning (day 0), mice underwent a 7 min protocol consisting of alternating 1 min silent periods and 30 s auditory stimulation periods (11-kHz tones delivered at 1 Hz). A 0.45 mA footshock was delivered at the end of each stimulation cycle, for a total of five tone–shock pairings. Freezing behavior was quantified during conditioning and recall sessions. Fear memory was evaluated by assessing generalized fear in a novel context (1 min, day 2), contextual fear in the conditioning chamber (1 min, day 3), and cued fear memory in response to the conditioned auditory stimulus (11-kHz, 1 Hz, 1 min) in a novel environment (day 4). Short-term memory formation was assessed using a protocol including conditioning (0 min), fear generalization (5 min), and cued fear recall (10 min), whereas long-term memory retention was evaluated over 5 days. For extinction experiments, mice underwent 15 consecutive 1 min tone presentations interleaved with 1 min silent intervals on day 4.
12. *Spatial memory* was assessed using the Morris water maze test, as previously described (Montarolo et al., 2013). The maze consisted of a circular pool (140 cm diameter, 40 cm height) filled with opaque water maintained at 25°C. A transparent escape platform (15 cm diameter) was positioned at a fixed location in the center of one quadrant and submerged 1 cm below the water surface. In 4 subsequent days, mice underwent four trials per day, with a 15-min inter-trial interval. Starting positions (north, south, east, or west) were randomized across trials. Mice were released facing the pool wall and allowed 90 s to locate the hidden platform. Animals failing to reach the platform within the given time were guided to it and allowed to remain there for 30 s. Between trials, mice were dried and housed in a cage containing paper towels. Escape latency was recorded and analyzed using EthoVision XT software (Noldus Information Technology, Wageningen, The Netherlands).

### Kainate-induced seizures

Seizures were induced in 3–4-month-old mice by intraperitoneal injection of the glutamatergic agonist kainic acid (30 mg/kg). Following administration, animals were individually housed in a Noldus PhenoTyper arena (30 × 30 cm) and video-recorded for 90 min using a high-definition frontal camera. Behavioral responses were analyzed offline using BORIS software (Friard and Gamba, 2016). Epileptic events were manually scored based on changes in exploratory activity, loss of posture, and overt seizure manifestations. Latency to seizure onset, number of seizures, and cumulative seizure duration were quantified for each animal.

### LFP recordings and analysis

To record extracellular field potentials (LFPs) in behaving mice, tungsten wire electrodes (Ø 75µm, AM Systems) were implanted under anesthesia (ip. ketamine-xylazine, 80mg/kg and 5 mg/kg, respectively). One group was implanted in the right and left M1 cortex (AP = +0.3 mm, ML = 1 mm from Bregma, and DV = -0.8 mm from the brain surface), with the reference electrode over the cerebellum. After 5 days of recovery, LFPs were recorded in freely moving mice while in their home-cage. A second cohort of animals was implanted in the right PFC (AP = +1.70 mm, ML = 0.4 mm from Bregma and DV = -2.3 mm from the brain surface) and right hippocampus (AP = -2 mm, ML = 2 mm from Bregma and DV = -1.5 mm from the brain surface), with the reference electrode over the left parietal cortex. In this group, 5 days after recovery, LFPs were recorded while mice were performing the Y-maze test. At the end of the recording, mice were sacrificed to verify electrode location using the Cresyl Violet staining (see Fig. 4D and Fig. 5C). In the first group, for each animal, five 2-second epochs were averaged during walking or resting periods. In the second group, five 2- second LFP epochs were selected and averaged. These epochs corresponded to the periods when mice were in the center of the maze, i.e. during the decision-making process. A customized Python script was used offline to measure coherence analysis (Marchiotto et al., 2025). Differences in coherence were measured according to the following frequency bands: delta (0.5-4 Hz), theta (4-12 Hz), beta (12-20), and gamma (20-30 Hz).

### Statistical Analysis

Statistical analyses were carried out with GraphPad Prism 9 (GraphPad software, Inc). The Shapiro-Wilk test was first applied to assess whether data followed a normal distribution. When normally distributed, unpaired Student’s t-test (to compare two groups) or Two-ways Anova (for multiple group comparisons) were used. Alternatively, when data were not normally distributed, the Mann–Whitney U-test was used. In all instances, P *<* 0.05 was considered as statistically significant. Histograms represent mean ± standard error (SE). Statistical differences were indicated with * P *<* 0.05, **P *<* 0.01, ***P *<* 0.001, ****P<0.0001. The list of the applied tests and the number of animals in each case are included in Suppl. Table 1. Source data are provided as a Source Data file (supplementary material).

## Results

### Oligodendroglial Cit-k deletion causes diffuse forebrain hypomyelination at juvenile stages

To investigate the contribution of oligodendroglial dysfunction to MCPH17 pathogenesis, we took advantage of the Sox10^Cre^;Cit-k^fl/fl^ mouse model, in which Cit-k is selectively deleted in oligodendroglial lineage cells, as validated in our previous study (Boda et al., 2022). At P14, mutant OPCs showed accumulation of DNA damage, increased apoptosis in the cerebral cortex, and senescence in the ventral forebrain, indicating that Cit-k loss disrupts oligodendroglial lineage homeostasis during early postnatal development. Based on these findings, we next investigated whether oligodendroglial Cit-k deletion affects CNS myelination and contributes to long-term neurological dysfunction.

First, we performed a gross neuroanatomical analysis of P14 Sox10^Cre^;Cit-k^fl/fl^ mice (Suppl. Fig. 1A-D). In contrast to constitutive Cit-k knockout mice (Di Cunto et al., 2000), mutant animals did not show microcephaly (i.e. no significant difference in forebrain hemispheric slice area and primary motor cortex (M1) thickness compared with controls; Suppl. Fig. 1A,C). However, mutant mice showed a trend toward ventriculomegaly (Suppl. Fig. 1B) and a significant reduction in corpus callosum (CC) thickness (Suppl. Fig. 1D), suggesting alterations in white matter.

To investigate whether oligodendroglial Cit-k deletion affected early CNS myelination, MBP expression was analyzed along the rostro-caudal axis of the forebrain at P14 (Fig. 1A). Sox10^Cre^;Cit- k^fl/fl^ mice displayed marked and diffuse hypomyelination involving both dorsal regions, including the cerebral cortex and CC, and ventral areas such as the striatum and hypothalamus (Fig. 1A,B). Consistent with these observations, western blot analysis revealed a strong reduction of the myelin proteins CNPase and MBP in both dorsal and ventral mutant forebrain, compared with age-matched controls (Fig. 1C,D). In contrast, levels of the axonal neurofilament marker SMI312 were unchanged (Fig. 1C,D), indicating that reduced myelination was not associated with axonal loss.

**Figure 1.**
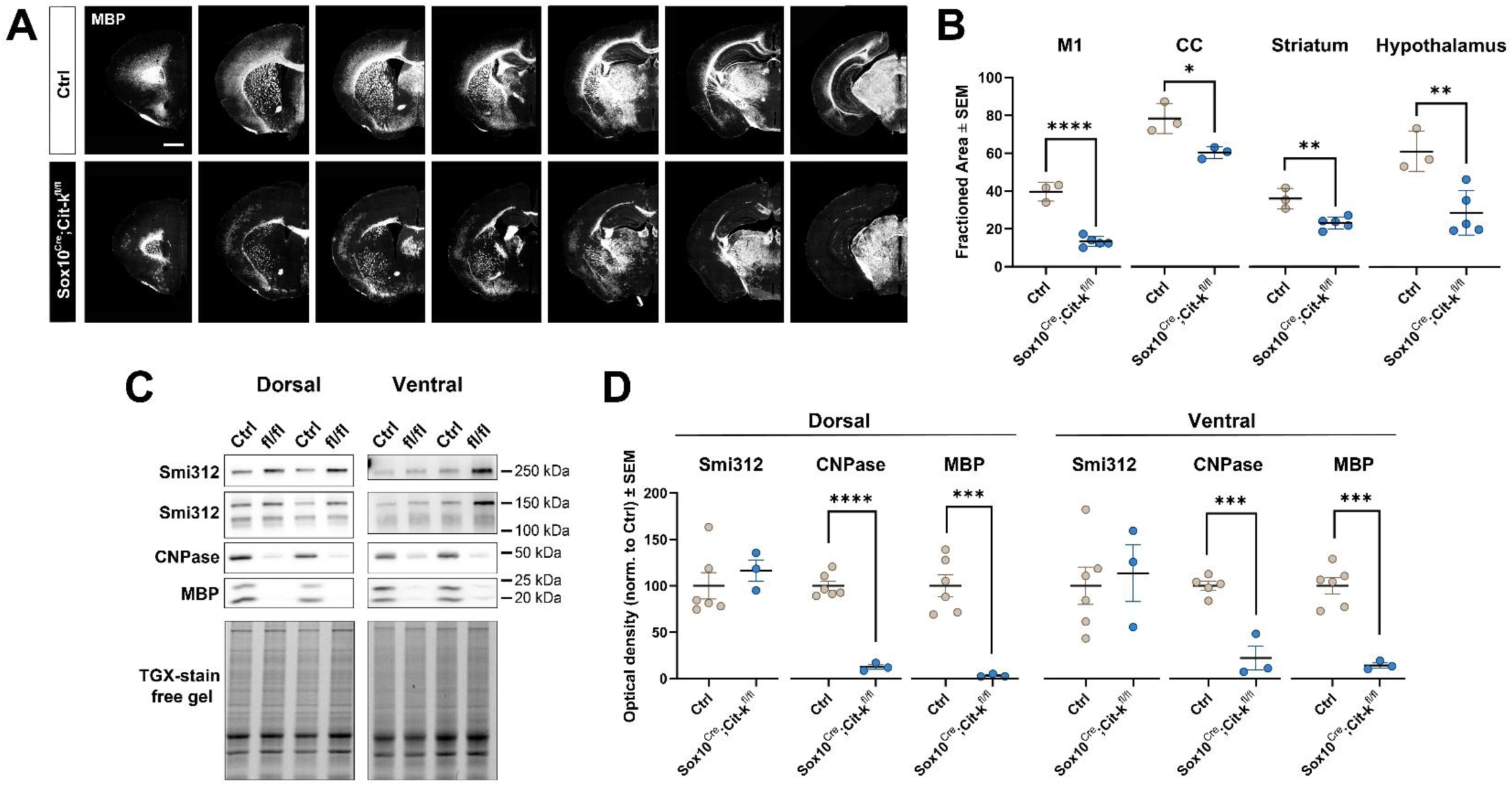
- Oligodendroglial Cit-k deletion causes diffuse forebrain hypomyelination in juvenile mice. **(A)** Representative images of MBP immunostaining in coronal forebrain sections from P14 control (Ctrl) and Sox10^Cre^;Cit-k^fl/fl^ mice at different rostro-caudal levels. Dorsal and ventral forebrain regions display markedly reduced MBP immunoreactivity in mutant animals. Scale bar: 500 µm. **(B)** Quantification of MBP-positive area fraction in the primary motor cortex (M1), corpus callosum (CC), striatum and hypothalamus. **(C)** Representative Western blots of the axonal neurofilament marker Smi312 and of the myelin proteins CNPase and MBP in dorsal and ventral forebrain lysates from P14 Ctrl and Sox10^Cre^;Cit-k^fl/fl^ mice. TGX stain-free gel imaging was used as loading control. **(D)** Quantification of Smi312, CNPase and MBP and protein levels in dorsal and ventral forebrain tissue normalized to total protein loading and to Ctrl levels. Each dot represents one mouse. Data are presented as mean ± SEM. Statistical significance was assessed by Student’s *t* test (see Suppl. Table 1). *P < 0.05, **P < 0.01, ***P < 0.001, **** P<0.0001. Source data are provided in the Source Data file.

Despite the marked forebrain hypomyelination, juvenile Sox10^Cre^;Cit-k^fl/fl^ mice did not show alterations in general motor development or vestibular function (Supplementary Fig. 2). Body weight, spontaneous motility, posture, and gait maturation were comparable between mutant and control littermates (Suppl. Fig. 2A–D). Similarly, vestibular reflexes assessed by negative geotaxis and surface righting tests (Suppl. Fig. 2E,F), as well as muscle strength evaluated by hindlimb suspension, were not significantly altered (Suppl. Fig. 2G).

To assess the development of early communicative behavior, we analyzed isolation-induced ultrasonic vocalizations (USVs). At P10, Sox10^Cre^;Cit-k^fl/f^ mice emitted a number of USVs comparable to that of control littermates (Suppl. Figs. 2H) and displayed no differences in the acoustic properties of individual calls (Suppl. Fig. 3A-I). Yet, mutant mice continued to emit spontaneous USVs following maternal/littermate isolation at P14, whereas control littermates had largely ceased vocalizing by this developmental stage (Suppl. Fig. 2H), suggesting a delayed maturation of the neural circuits underlying early communicative behavior.

### Oligodendroglial Cit-k loss causes long-lasting cortical myelin defects

We next investigated whether the myelination defects observed at juvenile stages persisted into adulthood. In adult Sox10^Cre^;Cit-k^fl/fl^ mice, myelination appeared largely recovered in major white matter tracts, including CC and fimbria, as well as in ventral telencephalic regions such as striatum, hypothalamus, and hippocampus (Fig. 2A,C). Accordingly, CC thickness and lateral ventricle size returned to levels comparable to those of control littermates (Suppl. Fig. 1F,H). In contrast, marked hypomyelination persisted throughout several cortical areas, including prefrontal/anterior cingulate (PFC/ACC), motor (M1) and sensory (S1, V1) cortices, as revealed by MBP immunostaining (Fig. 2A-C) and Gallyas myelin staining (Suppl. Fig. 4A,B). Western blot analysis of adult forebrain lysates confirmed the persistent reduction of CNPase and MBP protein levels in mutant mice, with the strongest decrease observed in dorsal forebrain regions (Fig. 2D,E). In the sensorimotor (M1-S1) cortex, cortical myelin displayed a characteristic uneven and patchy organization, with poorly myelinated regions alternating with completely unmyelinated areas in both deep (Fig. 2B, left panel) and superficial cortical layers (Fig. 2B, right panel). To investigate the cellular basis of such persistent cortical hypomyelination, we quantified oligodendroglial populations at different maturation stages in the sensorimotor cortex. While the density of NG2+ OPCs was comparable between genotypes (Fig. 3A,B), mutant mice showed a significant reduction in both BCAS1+ premyelinating (Fig. 3C,D) and CC1+ mature oligodendrocytes (Fig. 3E,F), indicating that persistent cortical hypomyelination is associated with a depletion of myelin-forming oligodendroglial cells rather than simply reflecting impaired myelin protein production.

**Figure 2.**
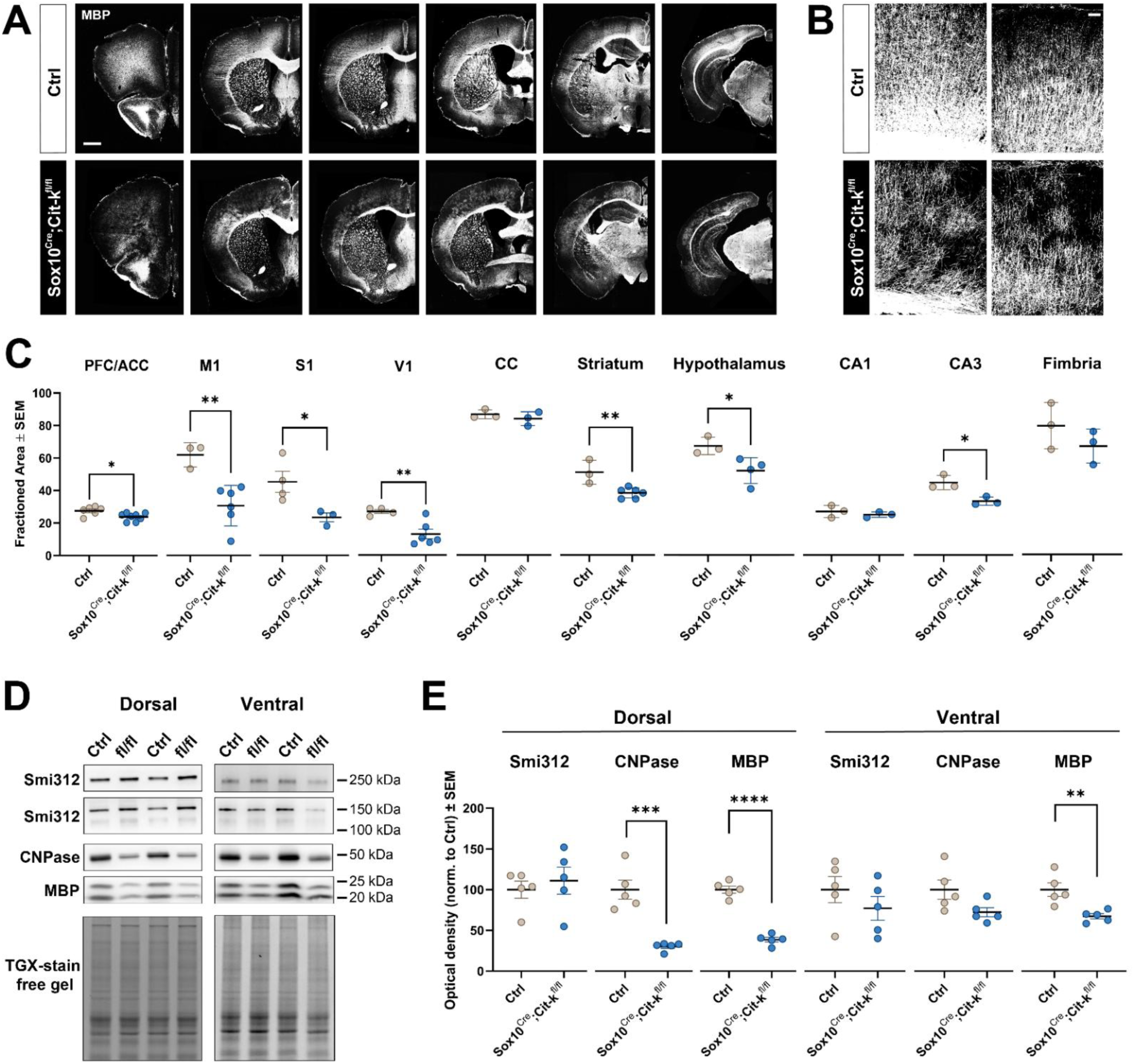
- Persistent cortical hypomyelination in adult Sox10^Cre^;Cit-k^fl/fl^ mice. **(A)** Representative images of MBP immunostaining in coronal forebrain sections from adult (3–5 months) control (Ctrl) and Sox10^Cre^;Cit-k^fl/fl^ mice at different rostro-caudal levels. Myelination is largely recovered in major white matter tracts and ventral forebrain regions, whereas reduced MBP immunoreactivity persists in several cortical areas. Scale bar: 500 µm. **(B)** High-magnification images of MBP immunostaining in the sensorimotor cortex (M1-S1). Mutant mice display a patchy myelination pattern characterized by poorly myelinated and unmyelinated regions in both deep (left panels) and superficial (right panels) cortical layers. Scale bar: 50 µm. **(C)** Quantification of MBP-positive area fraction in prefrontal/anterior cingulate cortex (PFC/ACC), primary motor cortex (M1), primary somatosensory cortex (S1), primary visual cortex (V1), corpus callosum (CC), striatum, hypothalamus, hippocampus (CA1, CA3), and fimbria. **(D)** Representative Western blots of Smi312, CNPase, and MBP in dorsal and ventral forebrain lysates from adult Ctrl and Sox10^Cre^;Cit-k^fl/fl^ mice. TGX stain- free gel imaging was used as loading control. **(E)** Quantification of Smi312, CNPase, and MBP protein levels in dorsal and ventral forebrain tissue normalized to total protein loading and to Ctrl levels. Each dot represents one mouse. Data are presented as mean ± SEM. Statistical significance was assessed by Student’s *t* test (see Suppl. Table 1). *P < 0.05, **P < 0.01, ***P < 0.001, **** P<0.0001. Source data are provided in the Source Data file.

**Figure 3.**
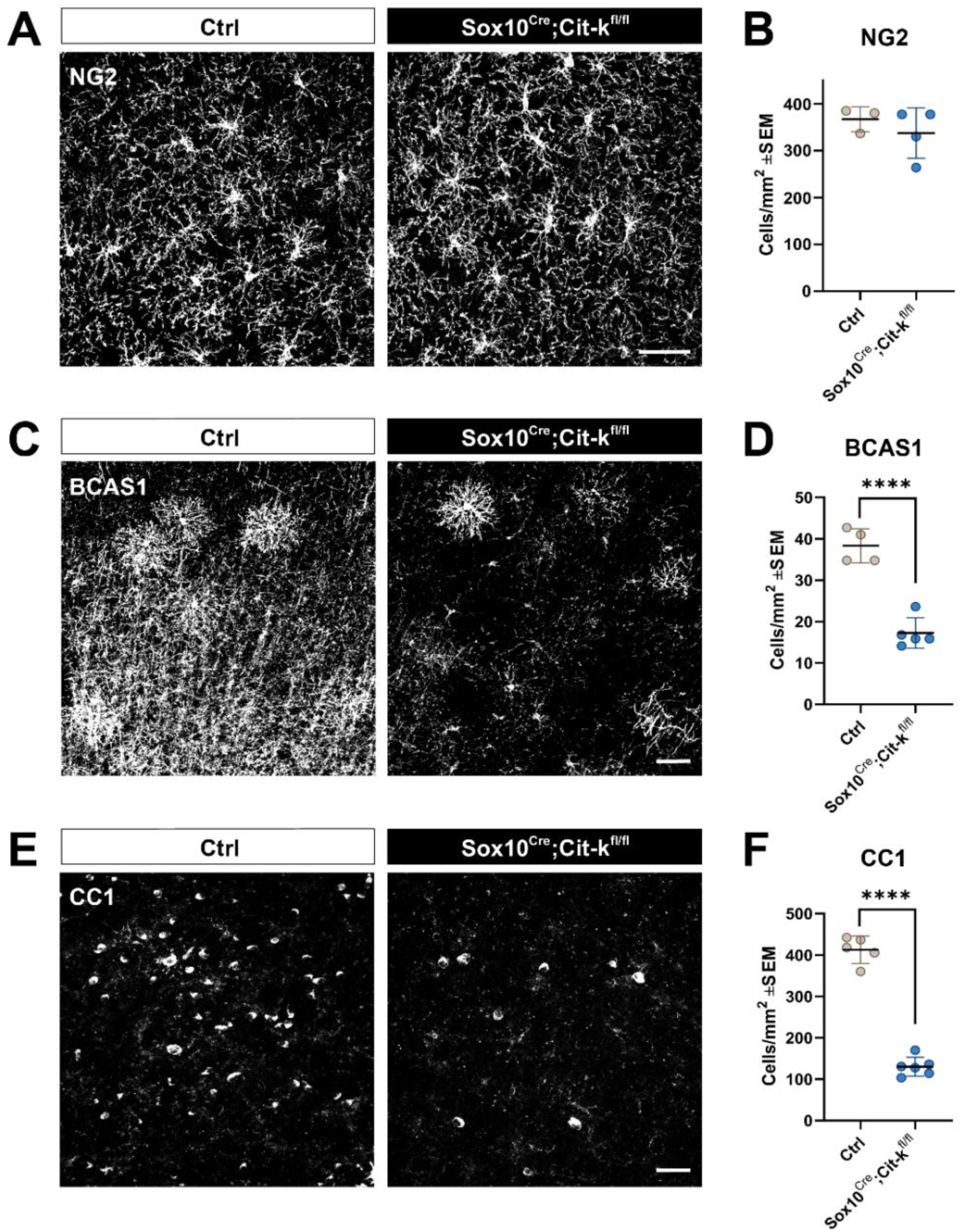
- Persistent cortical hypomyelination is associated with reduced numbers of premyelinating and mature oligodendrocytes. **(A,C,E)** Representative immunostainings of the sensorimotor cortex (M1-S1) from adult (3–5 months) control (Ctrl) and Sox10^Cre^;Cit-k^fl/fl^ mice showing NG2+ oligodendrocyte progenitor cells (OPCs; **A**), BCAS1+ premyelinating oligodendrocytes (**C**), and CC1+ mature oligodendrocytes (**E**). **(B,D,F)** Quantification of NG2+ OPC (**B**), BCAS1+ premyelinating oligodendrocyte (**D**), and CC1+ mature oligodendrocyte (**F**) cell density in the sensorimotor cortex. Scale bars: 50 µm. Each dot represents one mouse. Data are presented as mean ± SEM. Statistical significance was assessed by Student’s *t* test (see Suppl. Table 1). ****P < 0.0001. Source data are provided in the Source Data file.

### Subtle motor impairments in Sox10^Cre^;Cit-k^fl/fl^ mice are associated with reduced interhemispheric motor cortical synchrony

To determine whether persistent cortical hypomyelination was associated with functional abnormalities relevant to MCPH17, we next investigated sensory, motor, and cognitive functions in adult Sox10^Cre^;Cit-k^fl/fl^ mice. Mutant animals did not show major sensory impairments, as visual placing and auditory startle responses were comparable to those of controls (Suppl. Fig. 5A,B). Similarly, body weight, spontaneous locomotor activity, and hindlimb clasping behavior - commonly associated with spinal cord dysfunction - were not significantly altered in mutant animals (Suppl. Fig. 5C–E). Automated gait analysis using the CatWalk system revealed no major alterations in spontaneous locomotion. All principal gait parameters, e.g. walking speed, stride length and regularity, and base of support, were comparable between genotypes, with only minor differences in print position (i.e. the distance between the placement of the hindpaw and the preceding ipsilateral forepaw) and lateral support (relative duration of simultaneous contact of the left or right limb pairs; Suppl. Fig. 5F). Gross motor coordination and motor learning were also preserved, as assessed by accelerating rotarod performance across 3 consecutive days (3 trials/day), during which mutant and control mice showed comparable latency to fall and learning curves (Suppl. Fig. 5G).

Yet, movement initiation appeared mildly but significantly impaired in mutant mice, as evidenced by the increased latency to release an externally imposed posture in the bar test (Fig. 4A), consistent with previous observations in other mutants of oligodendroglial genes (Hagemeyer et al., 2012). Fine motor coordination was also impaired in mutant mice, as indicated by the increased number of lateral slips and the tendency to require longer time to traverse the beam in the beam-walking test (Fig. 4B,C).

**Figure 4.**
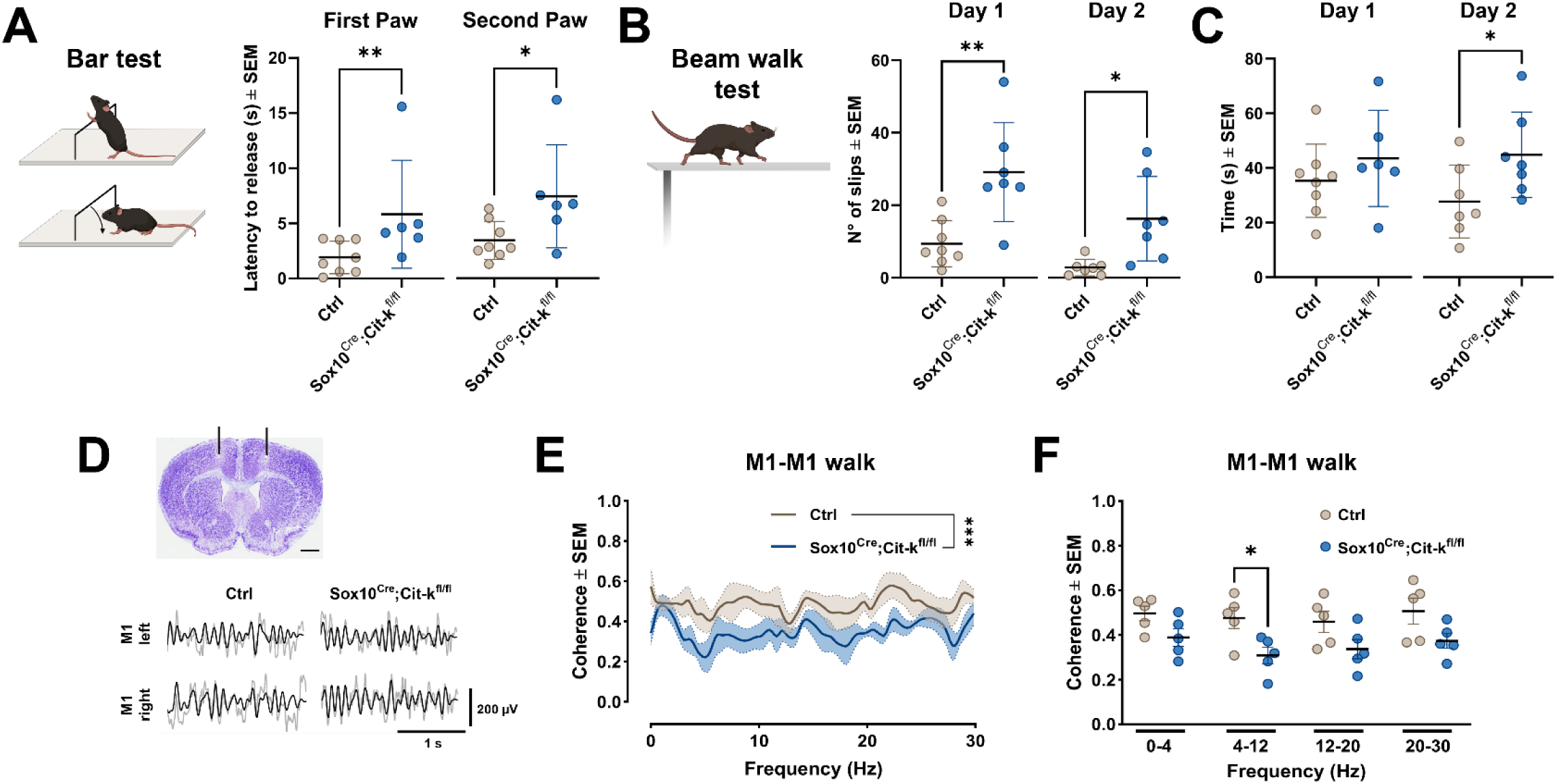
- Subtle motor deficits are associated with reduced interhemispheric M1-M1 synchrony. **(A)** Bar test performance in adult (3–5 months) control (Ctrl) and Sox10^Cre^;Cit-k^fl/fl^ mice. Latency to release the first and second forepaw from an externally imposed posture was used as an index of movement initiation ability. **(B,C)** Beam- walking test performance. **(B)** Number of hindlimb lateral slips during beam traversal. **(C)** Time required to traverse the beam. **(D)** Assessment of interhemispheric motor cortical synchrony during spontaneous walking. Upper panel: representative Cresyl Violet-stained coronal section showing electrode placement in the left and right primary motor cortices (M1). Scale bar: 1 mm. Lower panel: representative LFP traces recorded in the right/left M1 of control (Ctrl) and Sox10^Cre^;Cit-k^fl/fl^ mice. Raw 1–30 Hz filtered traces are shown in gray. 4–12 Hz filtered traces are overlaid and shown in black. **(E)** Mean coherence spectrum between M1 LFPs recorded simultaneously from the two hemispheres during walking. **(F)** Mean M1-M1 coherence values in delta (0.5–4 Hz), theta (4–12 Hz), beta (12–20 Hz), and gamma (20–30 Hz) frequency bands during walking. Each dot represents one mouse. Data are presented as mean ± SEM. Statistical significance was assessed by Student’s *t* test, Mann–Whitney *U* test, or Two-way ANOVA, as appropriate (see Suppl. Table 1). *P < 0.05, **P < 0.01; ***P < 0.001. Source data are provided in the Source Data file. Illustrations in A,B were created with BioRender.com.

The absence of detectable hypomyelination in the cerebellum and spinal cord (Suppl. Fig.4A) suggests that the subtle motor deficits observed in mutant mice are unlikely to originate from these structures. Rather, they are consistent with the persistent hypomyelination affecting sensorimotor cortical regions (see above) and may reflect an altered interhemispheric connectivity between motor cortical areas. To test this possibility, we simultaneously recorded extracellular LFPs from the two M1 cortices and quantified interhemispheric synchrony during spontaneous walking (Fig. 4D). Increased coherence between the two regions is generally interpreted as a measure of functional coupling (Marchiotto et al., 2025; Jeong et al., 2021), whereas reduced interhemispheric synchronization is commonly associated with impaired bilateral motor coordination (Lemke et al., 2019). A general reduction in interhemispheric M1-M1 coherence across the analyzed frequency spectrum was detected in mutant mice compared with controls, indicating decreased functional coupling between the two motor cortices during walking. This effect reached statistical significance within the 4–12 Hz (theta) frequency range (Fig. 7E,F). Notably, theta-frequency oscillations in rodents have been linked to locomotion-related motor network dynamics (Noga et al., 2017; Vanni et al., 2017), suggesting that reduced M1–M1 synchronization may contribute to the motor deficits observed in mutant mice. In contrast, M1–M1 coherence was not different between genotypes when animals were resting (Suppl. Fig. 6A,B), indicating that reduced interhemispheric coupling becomes evident primarily during locomotion.

### Memory deficits in Sox10^Cre^;Cit-k^fl/fl^ mice are associated with disrupted cortico-hippocampal connectivity

Given that cognitive impairment is a major feature of MCPH17, we next examined the impact of oligodendroglial Cit-k loss on cognitive function. Spatial working memory was assessed using the spontaneous alternation Y-maze task. Mutant mice exhibited a reduced spontaneous alternation rate, reflecting a lower number of correct arm choices, while the total number of arm entries was unchanged between genotypes (Fig. 5A). These findings indicate impaired spatial working memory in the absence of major alterations in exploratory behavior. Spatial working memory critically relies on the crosstalk between the PFC and hippocampus, which are coordinated by synchronized oscillatory activity, particularly within the theta frequency range, at the choice point (Benchenane et al., 2010). To determine whether the working memory deficits observed in mutant mice were associated with altered cortico-hippocampal connectivity, we simultaneously recorded extracellular LFPs from the PFC and hippocampus during the decision phase of the Y-maze task, corresponding to the 2 seconds preceding arm entry (Fig. 5B,C). Control mice displayed a marked increase in PFC– hippocampal synchrony within the 4–12 Hz range during arm choice, whereas this coherence peak was abolished in mutant animals (Fig. 5C,D). These findings indicate that impaired working memory performance in Sox10^Cre^;Cit-k^fl/fl^ mice is associated with defective cortico-hippocampal functional coupling.

**Figure 5.**
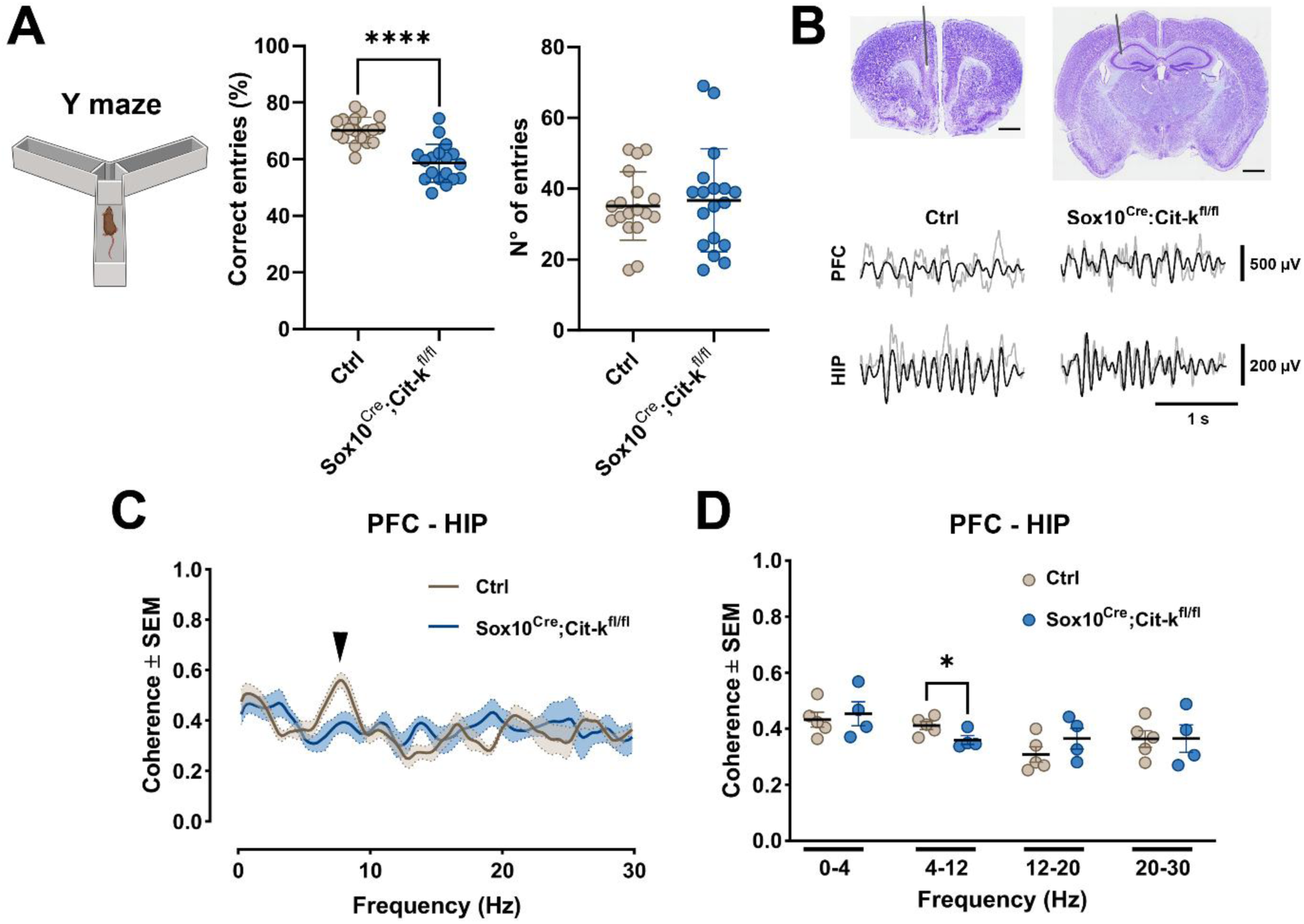
- Working memory deficits are associated with reduced theta-range cortico-hippocampal synchrony. **(A)** Left panel: spontaneous alternation performance in the Y-maze, expressed as the percentage of correct arm entries. Right panel: total number of arm entries during the Y-maze task. **(B)** Assessment of cortico-hippocampal synchrony during the decision phase of the Y-maze task. Upper panel: representative Cresyl Violet-stained coronal sections showing electrode placement in the PFC and hippocampus. Scale bar: 1 mm. Lower panel: representative LFP traces recorded in the prefrontal cortex (PFC) and hippocampus (HIP) of control (Ctrl) and Sox10^Cre^;Cit-k^fl/fl^ mice. Raw 1–30 Hz filtered traces are shown in gray. 4–12 Hz filtered traces are overlaid and shown in black. **(C)** Mean coherence spectrum between PFC and HIP LFPs recorded during the 2 s preceding arm entry. **(D)** Mean PFC–HIP coherence values in delta (0.5–4 Hz), theta (4–12 Hz), beta (12–20 Hz), and gamma (20–30 Hz) frequency bands during the decision phase of the task. Each dot represents one mouse. Data are presented as mean ± SEM. Statistical significance was assessed by Student’s t test or Two-way ANOVA, as appropriate (see Supplementary Table 1). *P < 0.05; ****P < 0.0001. Source data are provided in the Source Data file. Illustration in A was created with BioRender.com.

We next extended our analysis to other cognitive domains. In the novel object recognition (NOR) test, using a 2.5 h intertrial interval, Sox10^Cre^;Cit-k^fl/fl^ mice exhibited a significant impairment in short-term recognition memory, as evidenced by a reduced preference for the novel object compared with control mice (Fig. 6A,B). Notably, recognition memory has also been associated with functional interactions between the hippocampus and PFC during novelty discrimination (Wang et al., 2021). Next, we examined emotional learning, short-term and long-term memory by investigating contextual and auditory fear memory (Fig. 6C and Suppl Fig. 7A). Mutant and control mice displayed comparable freezing responses during conditioning and in the immediate post-conditioning period (Suppl. Fig. 7A–C), indicating normal expression of the freezing response. Consistent with this, no genotype-dependent differences were observed during short-term auditory fear recall, assessed 10 min after conditioning, when both groups spent approximately 80% of the time freezing in response to the conditioned auditory stimulus (Fig. 6D). Similarly, neither fear generalization, evaluated in a novel environment 5 min or 2 days after conditioning (Suppl. Fig. 7D,E), nor contextual fear memory, assessed 3 days after conditioning (Fig. 6E,F), differed between genotypes. The latter results suggest that a simple form of hippocampal-dependent memory, such as contextual fear memory, is preserved in these mutant mice. In contrast, mutant mice exhibited a significantly reduced and more variable (coefficient of variation, CV: 49.13 *vs* 17.05) freezing response during long-term auditory recall performed 4 days after conditioning (Fig. 6G), indicating a selective impairment in the long-term auditory fear memory retention. This deficit was associated with relatively mild myelin alterations in the primary (Te1) and temporal association (Te2) auditory cortices and absence of detectable hypomyelination in the amygdala (Fig. 2A and Suppl. Fig. 7F), which are critically involved in the consolidation, storage and retrieval of auditory fear memories (Cambiaghi et al., 2016; Concina et al., 2024). Memory extinction appeared preserved, as mutant and control mice displayed a similar progressive decline in freezing responses during repeated presentations of the conditioned stimulus, despite the lower initial freezing levels observed in mutant animals (Fig. 6H).

**Figure 6.**
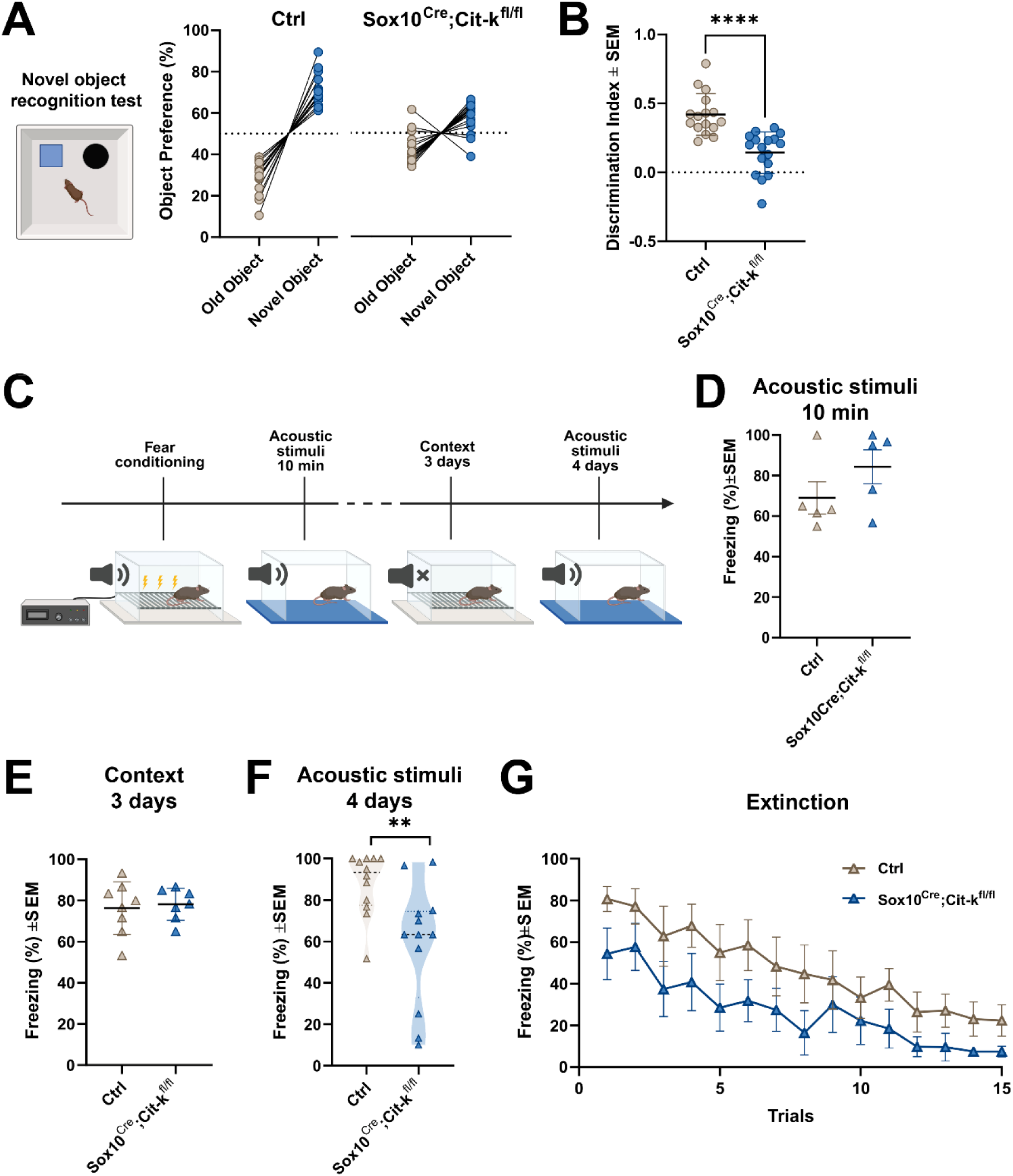
- Oligodendroglial Cit-k loss impairs recognition memory and long-term auditory fear memory. **(A)** Novel object recognition (NOR) performance, expressed as preference for the novel object. **(B)** NOR discrimination index. **(C)** Experimental design of the auditory fear-conditioning paradigm. Mice underwent conditioning on day 0 followed by short-term cued fear recall (10 min), contextual fear recall (day 3), long-term cued fear recall (day 4), and extinction testing. **(D)** Short-term cued fear recall assessed 10 min after conditioning and expressed as percentage of time spent freezing during presentation of the conditioned auditory stimulus. **(E)** Contextual fear memory assessed 3 days after conditioning. **(F)** Long-term cued fear memory assessed 4 days after conditioning. Violin plots show the distribution and variability (i.e. median, quartiles and frequency distribution density of data) of freezing responses in each genotype. **(G)** Fear extinction assessed during 15 repeated presentations of the conditioned auditory stimulus on day 4. Each dot represents one mouse, except in (G), where data are presented as mean ± SEM. Statistical significance was assessed by Student’s *t* test, Mann–Whitney *U* test, or Two-way ANOVA, as appropriate (see Suppl. Table 1). **P < 0.01; ****P < 0.0001. Source data are provided in the Source Data file. Illustrations in A,C were created with BioRender.com.

Finally, spatial learning and memory were assessed using the Morris water maze. Overall, Sox10^Cre^;Cit-k^fl/fl^ mice displayed mean escape latencies comparable to those of control mice (Suppl. Fig. 7G). However, mutant mice exhibited higher inter-individual variability in performance across training sessions (CV: 90.75 *vs* 49.9 on the 4^th^ day). While some mutant animals learned the task similarly to controls, others failed to show the expected reduction in escape latency despite repeated training. Thus, spatial learning is not uniformly impaired following oligodendroglial Cit-k loss, but rather becomes less consistent across individuals, possibly reflecting differences in compensatory strategies or in the recruitment of cortical and hippocampal networks supporting spatial navigation (see Discussion).

Collectively, these findings indicate that oligodendroglial Cit-k loss impairs multiple cognitive domains relying on cortical and hippocampal circuits and identify defective cortico-hippocampal synchronization as a potential network mechanism underlying these abnormalities.

### Oligodendroglial Cit-k deletion increases susceptibility to kainate-induced seizures

Given the occurrence of seizures in a subset of MCPH17 patients, we examined seizure susceptibility in Sox10^Cre^;Cit-k^fl/fl^mice. Although no spontaneous epileptic phenotype was detected at juvenile and adult ages, mutant mice showed a pronounced increase in susceptibility to kainate-induced seizures. Following administration of kainic acid, mutants displayed a shorter latency to seizure onset (approximately 16 *vs* 50 min), a greater number of seizures, and a tendency toward longer seizure duration during the 90 min observation period (Fig 7A-C). These findings indicate that oligodendroglial *Cit-k* loss lowers the threshold for seizure generation, linking connectivity defects to enhanced vulnerability to epileptogenic perturbations.

**Figure 7.**
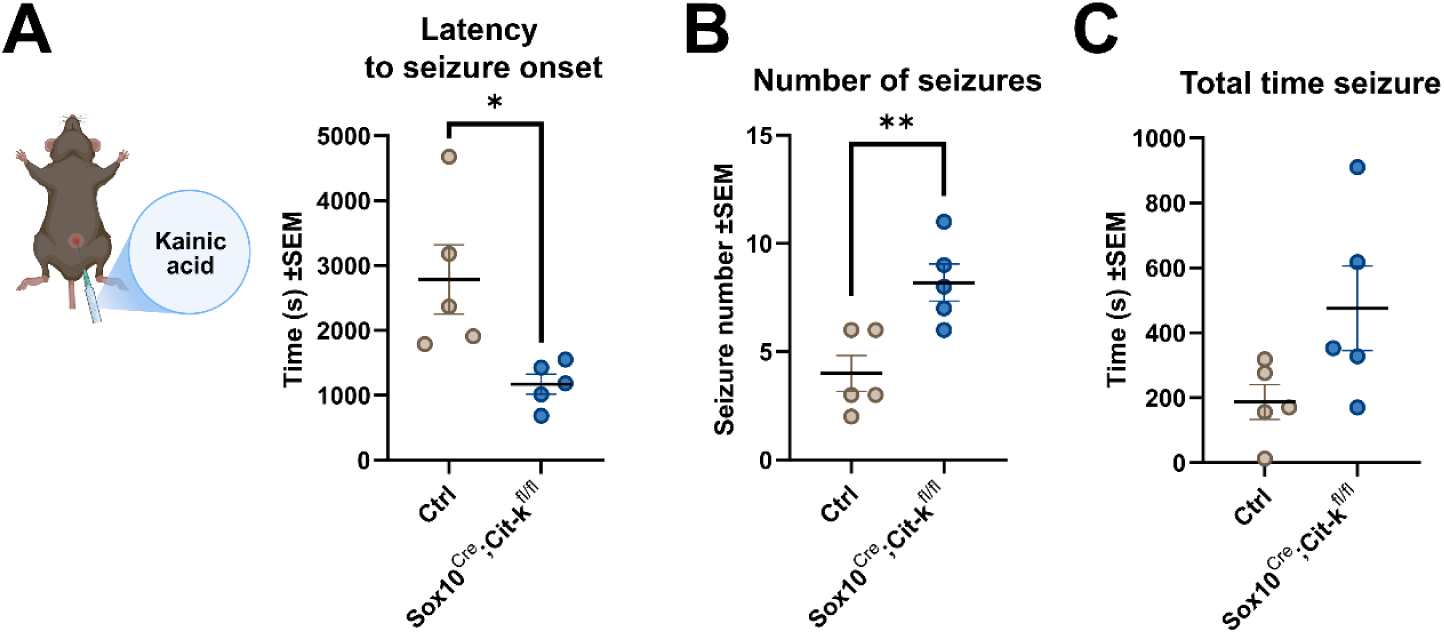
- Oligodendroglial Cit-k loss increases susceptibility to kainate-induced seizures. Adult (3-5 months) control (Ctrl) and Sox10^Cre^;Cit-k^fl/fl^ mice were injected with kainic acid and monitored for seizure activity during a 90 min observation period. **(A)** Latency to seizure onset, measured as the time to first loss of posture following kainic acid administration. **(B)** Total number of seizures observed during the recording period. **(C)** Cumulative seizure duration during the recording period. Each dot represents one mouse. Data are presented as mean ± SEM. Statistical significance was assessed by Student’s *t* test or Mann–Whitney *U* test (see Suppl.Table 1). *P < 0.05, **P < 0.01. Source data are provided in the Source Data file. Illustration in A was created with BioRender.com.

## Discussion

The present study demonstrates that selective loss of Cit-k in oligodendroglial lineage cells is sufficient to produce long-lasting neurological abnormalities independently of primary neuronal defects. By combining anatomical, behavioral, electrophysiological, and seizure susceptibility analyses, we show that oligodendroglial dysfunction associated with Cit-k loss can impair *per se* motor and cognitive functions, alter cortico-cortical and cortico-hippocampal connectivity, and increase seizure vulnerability. Importantly, these functional abnormalities occurred in the absence of overt microcephaly or major neuroanatomical defects. This dissociation indicates that neurological dysfunction in MCPH17 cannot be explained solely by reduced brain size and highlights the contribution of non-neuronal mechanisms to disease outcome.

MCPH has traditionally been regarded as a prototypical neuronal disorder, with disease manifestations largely attributed to impaired neural progenitor proliferation and the consequent reduction of neuronal populations. In constitutive Cit-k knockout mice, severe microcephaly, extensive neuronal loss, and early lethality (Di Cunto et al., 2000) have reinforced this interpretation. However, our previous work demonstrated that Cit-k also plays a cell-autonomous role in oligodendroglial lineage cells, where its loss induces DNA damage accumulation, apoptosis, and senescence (Boda et al., 2022). The present study extends these observations by showing that oligodendroglial Cit-k deletion is sufficient to generate persistent neurological dysfunctions in the absence of primary neuronal defects. Together, these findings indicate that oligodendroglial abnormalities are not simply secondary consequences of impaired neurogenesis but contribute directly to disease pathogenesis and neurological outcome.

Hypomyelination represents the most obvious candidate mechanism linking oligodendroglial dysfunction to neurological outcome. A key finding of this study is that the severe and diffuse hypomyelination observed during development was followed by only a partial recovery. While myelination largely normalized in major white matter tracts and several subcortical regions, hypomyelination persisted across the cerebral cortex and was accompanied by long-lasting motor and cognitive impairments, functions that critically depend on cortical processing and coordinated communication between cortical and subcortical regions.

The mechanisms underlying this regional vulnerability remain unclear. Cortical oligodendroglial populations may display a greater dependence on Cit-k function, consistent with the developmental and regional heterogeneity of forebrain OPCs (Boda et al., 2022; Foerster et al., 2024; Foerster et al., 2019). Alternatively, the delayed and protracted maturation of cortical myelin may render cortical oligodendroglial cells particularly vulnerable to perturbations affecting lineage progression and maturation (de Faria et al., 2021). Interestingly, hypomyelination was not uniformly distributed across cortical regions. Sensorimotor cortices, which are among the most heavily myelinated cortical areas, exhibited the most pronounced deficits (-50% of MBP+ area), whereas less myelinated regions, such as the PFC/ACC (-14%) or Te1-Te2 (-20%) cortices, were less affected. This pattern suggests that the consequences of Cit-k loss may depend, at least in part, on regional differences in myelination demand and oligodendroglial biology.

Importantly, persistent cortical hypomyelination was associated with a marked reduction of both premyelinating and mature oligodendrocytes, whereas OPC density largely recovered by adulthood. These findings indicate that the adult phenotype is not simply the consequence of a lasting depletion of progenitor cells. Rather, they suggest that Cit-k function, including its roles in cytoskeletal dynamics and genome integrity maintenance (Di Cunto et al., 2000; Bianchi et al., 2017; Iegiani et al., 2025), may be particularly important for the generation and/or maintenance of myelin-forming oligodendrocytes. Whether and how Cit-k directly regulates specific transitions along the oligodendroglial lineage or is required for the long-term survival of differentiated oligodendrocytes remains to be determined.

In adult mice, cortical hypomyelination was associated with functional connectivity alterations in both motor and cognitive networks. Reduced M1-M1 synchronization emerged during locomotion and was accompanied by subtle impairments in movement initiation and motor accuracy, whereas working memory deficits were associated with reduced PFC-hippocampal coupling during the decision phase of the Y-maze task. In both cases, the effect was most evident within the theta frequency range, an oscillatory band known to support the corresponding behaviors (Noga et al., 2017; Vanni et al., 2017; Benchenane et al., 2010). These findings suggest that oligodendroglial dysfunction impairs the coordination of neural activity across distributed brain regions during behavior.

Interestingly, not all cognitive functions were affected to the same extent. While mutant mice displayed clear deficits in working memory, recognition memory, and long-term auditory fear memory, contextual fear memory and spatial learning were largely preserved at the group level. Nevertheless, in the Morris water maze, mutant animals showed greater inter-individual variability, with some performing comparably to controls and others showing limited learning. These findings suggest that oligodendroglial dysfunction may reduce the efficiency and reliability of the neural networks supporting cognition rather than causing a uniform impairment across individuals.

Given the critical role of myelin in regulating action potential conduction and in supporting efficient communication between brain regions, altered synchronization may represent an important mechanism linking oligodendroglial pathology to network and behavioral dysfunction. However, our findings do not allow us to attribute these phenotypes exclusively to impaired myelin deposition.

Notably, some behavioral alterations were observed even in the absence of marked hypomyelination in the regions classically associated with the affected function. For example, impaired long-term auditory fear memory occurred despite relatively mild hypomyelination in Te1 and Te2 and the absence of detectable myelin alterations in amygdala (Cambiaghi et al., 2016; Concina et al., 2022; Concina et al., 2024), suggesting that even subtle myelin abnormalities may be sufficient to perturb neural networks. At the same time, the persistent reduction of premyelinating and mature oligodendrocytes observed in the adult cortex raises the possibility that additional oligodendroglial functions also contribute to the phenotype. Beyond producing myelin, oligodendrocytes contribute to neural circuit development and function through metabolic support, regulation of neuronal excitability, axonal remodeling, and synapse engulfment (Schirmer et al., 2018; Simons et al., 2024; Buchanan et al., 2023). The loss of these populations may therefore contribute to the observed behavioral and connectivity abnormalities through mechanisms that extend beyond impaired myelin deposition. Future studies will be required to disentangle the relative contribution of myelin-dependent and myelin-independent oligodendroglial functions to disease pathogenesis.

The increased susceptibility of Sox10^Cre^;Cit-k^fl/fl^ mice to kainate-induced seizures provides further evidence that oligodendroglial dysfunction compromises network stability. Although spontaneous seizures were not observed, mutant animals displayed a reduced latency to seizure onset and an increased seizure burden following kainate administration, indicating a lower threshold for seizure generation. This finding is consistent with the occurrence of epileptic manifestations in a subset of MCPH17 patients and suggests that oligodendroglial alterations may contribute directly to seizure susceptibility. The mechanisms underlying this phenotype remain unclear. One possibility is that hypomyelination and/or oligodendroglial dysfunction alters the organization and communication of long-range brain networks, thereby increasing their vulnerability to epileptogenic perturbations. In addition, oligodendroglia and hypomyelination may affect local circuit function, including the maturation and activity of parvalbumin-positive interneurons (Stedehouder et al., 2017; Benamer et al., 2020; Dubey et al., 2022; Plaisier et al., 2025), which play a key role in cortical synchronization and seizure control (Sohal et al., 2009; Trevelyan et al., 2006).

Beyond their mechanistic implications, our findings suggest that oligodendroglial dysfunction may represent a therapeutic target in MCPH17. Unlike the primary neurogenic defects underlying congenital microcephaly, oligodendroglial abnormalities develop largely during postnatal stages (Khelfaoui et al., 2024), potentially offering a broader window for intervention. Restoring oligodendrocyte function or myelination could therefore ameliorate connectivity deficits, motor and cognitive impairment, and seizure susceptibility even after developmental neuronal deficits have occurred.

In conclusion, our findings show that selective oligodendroglial Cit-k loss is sufficient to cause persistent defects in myelination, brain connectivity, cognition, and seizure susceptibility independently of primary neuronal abnormalities. These results identify oligodendroglial dysfunction as an important contributor to MCPH17 pathogenesis and further support a role for myelin/oligodendroglial abnormalities in the emergence of network and behavioral dysfunction across neurodevelopmental disorders.

## Supporting information

Souce Data File

---

**Supplementary Figure 1.**
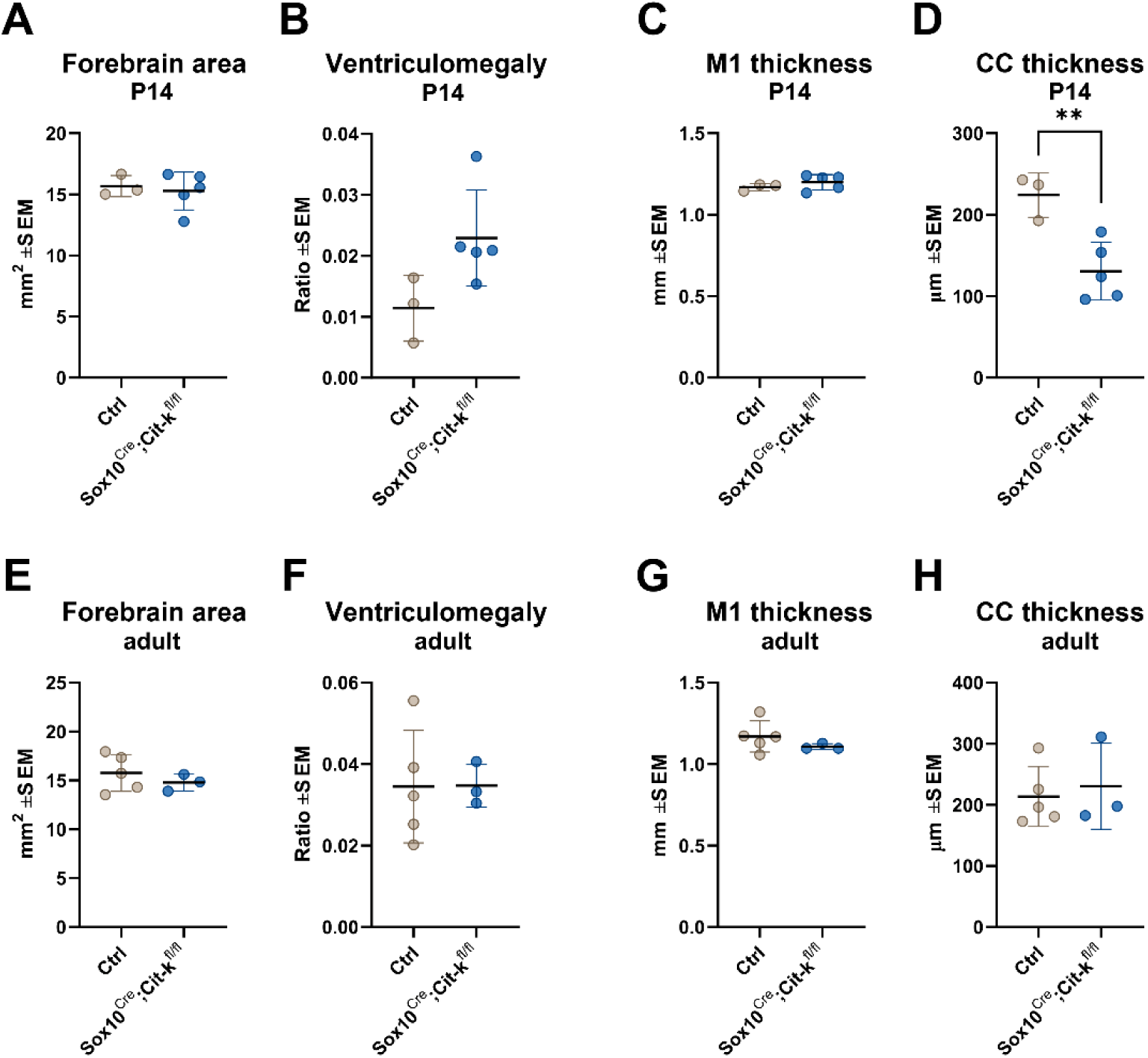
– Gross neuroanatomical development is largely preserved in Sox10^Cre^;Cit- k^fl/fl^ mice. Quantification of forebrain morphological parameters in juvenile (P14; **A–D**) and adult (3–5 months; **E–H**) mice. **(A,E)** Forebrain hemispheric area measured at the level of the lateral ventricles. **(B,F)** Ventriculomegaly index, calculated as the ratio between lateral ventricular area and total hemispheric area. **(C,G)** Thickness of the primary motor cortex (M1). **(D,H)** Corpus callosum (CC) thickness. Each dot represents one mouse. Statistical significance was assessed by Student’s *t* test or Mann–Whitney *U* test (see Suppl. Table 1). **P < 0.01. Source data are provided in the Source Data file.

**Supplementary Figure 2.**
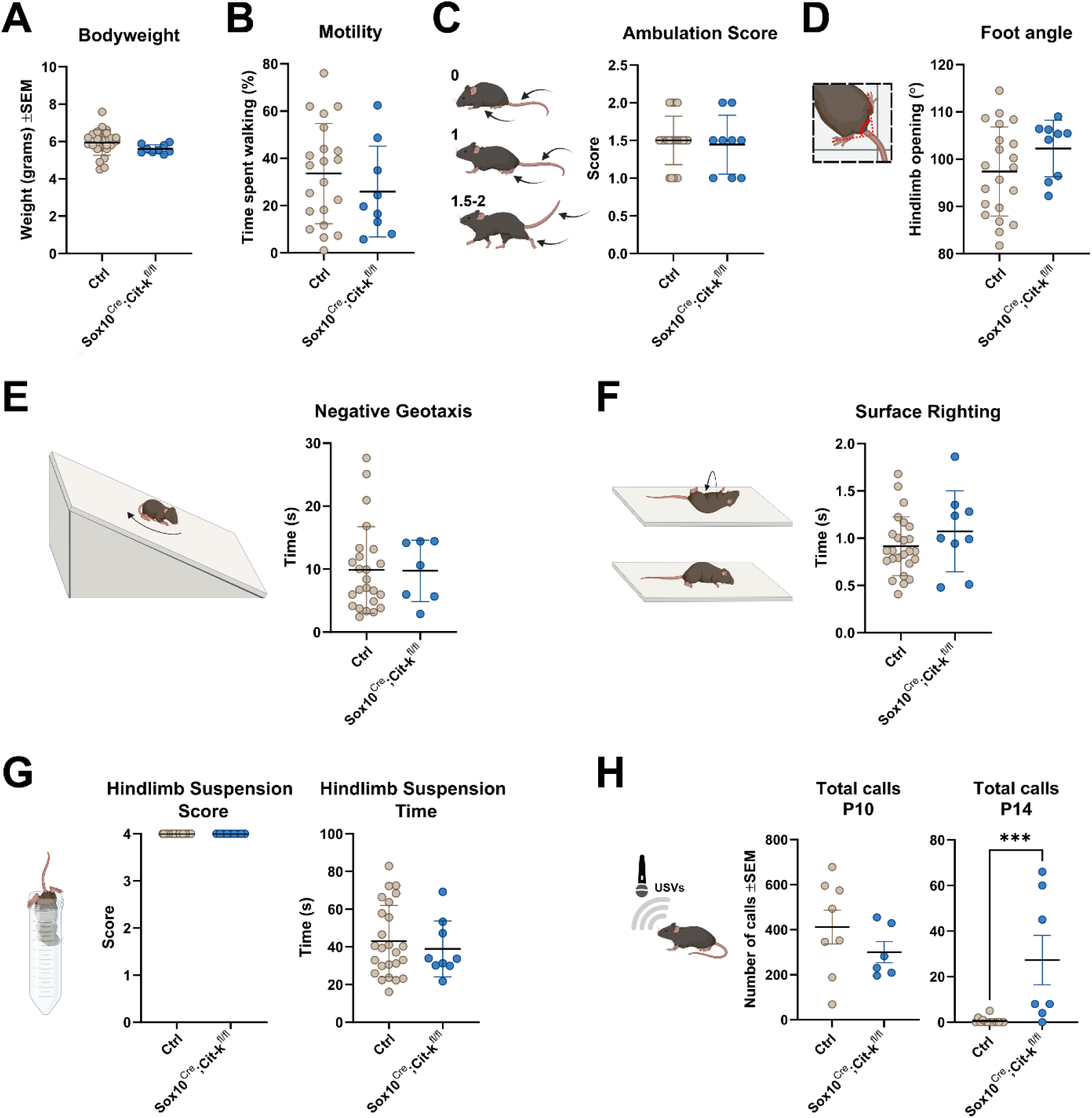
- Preserved motor and vestibular development but persistent ultrasonic vocalizations in juvenile Sox10^Cre^;Cit-k^fl/fl^ mice. Behavioral characterization of juvenile (P14) control (Ctrl) and Sox10^Cre^;Cit-k^fl/fl^ mice. **(A)** Body weight. **(B)** Spontaneous motility, expressed as the percentage of time spent moving in an open-field arena. **(C)** Ambulation score, as a measure of gait maturation. **(D)** Hindlimb foot angle, as an index of postural control. **(E)** Negative geotaxis performance (time to complete the task). **(F)** Surface righting reflex time. **(G)** Hindlimb suspension test. Left, hindlimb posture score. Right, latency to fall. **(H)** Number of isolation-induced ultrasonic vocalizations (USVs) emitted at P10 and P14. Each dot represents one mouse. Data are presented as mean ± SEM. Statistical significance was assessed by Student’s *t* test, or Mann–Whitney *U* test, as appropriate (see Suppl. Table 1). *P < 0.05, **P < 0.01, ***P<0.001. Source data are provided in the Source Data file. Illustrations were created with BioRender.com.

**Supplementary Figure 3.**
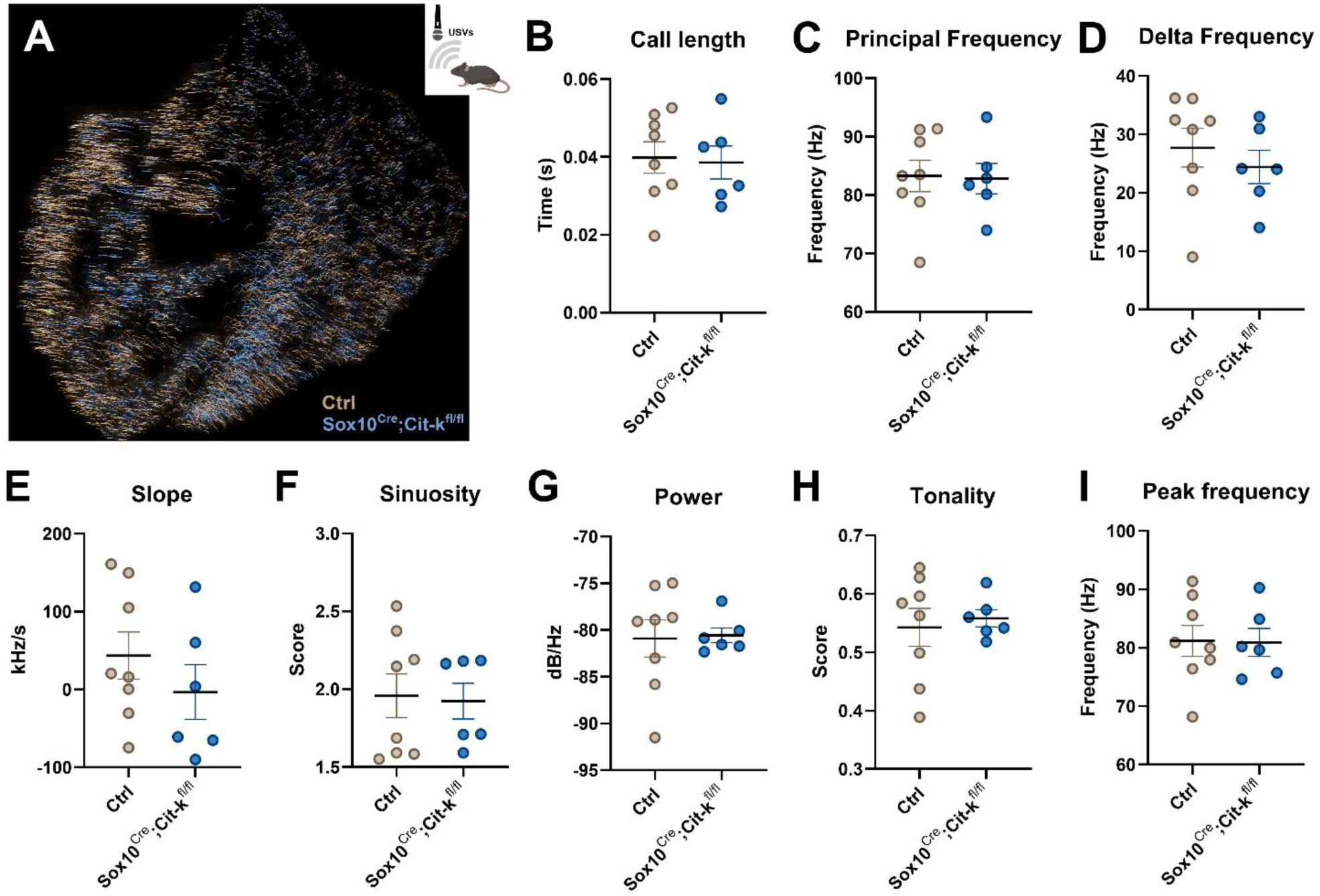
- Acoustic features of isolation-induced ultrasonic vocalizations are preserved in juvenile Sox10^Cre^;Cit-k^fl/fl^ mice. DeepSqueak-based analysis of isolation-induced ultrasonic vocalizations (USVs) recorded from juvenile (P10) mice. **(A)** Ctrl and Sox10^Cre^;Cit-k^fl/fl^ USV traces largely overlaps in the 2D visualization after UMAP embedding and clustering for acoustic features. Quantification of call duration **(B)**, principal frequency **(C)**, delta frequency **(D)**, slope **(E)**, sinuosity **(F)**, power **(G)**, tonality **(H)**, and peak frequency **(I)**. Each dot in B-I represents one mouse. Data are presented as mean ± SEM. Statistical significance was assessed by Student’s *t* test (see Suppl. Table 1). Source data are provided in the Source Data file. Illustration in A was created with BioRender.com.

**Supplementary Figure 4.**
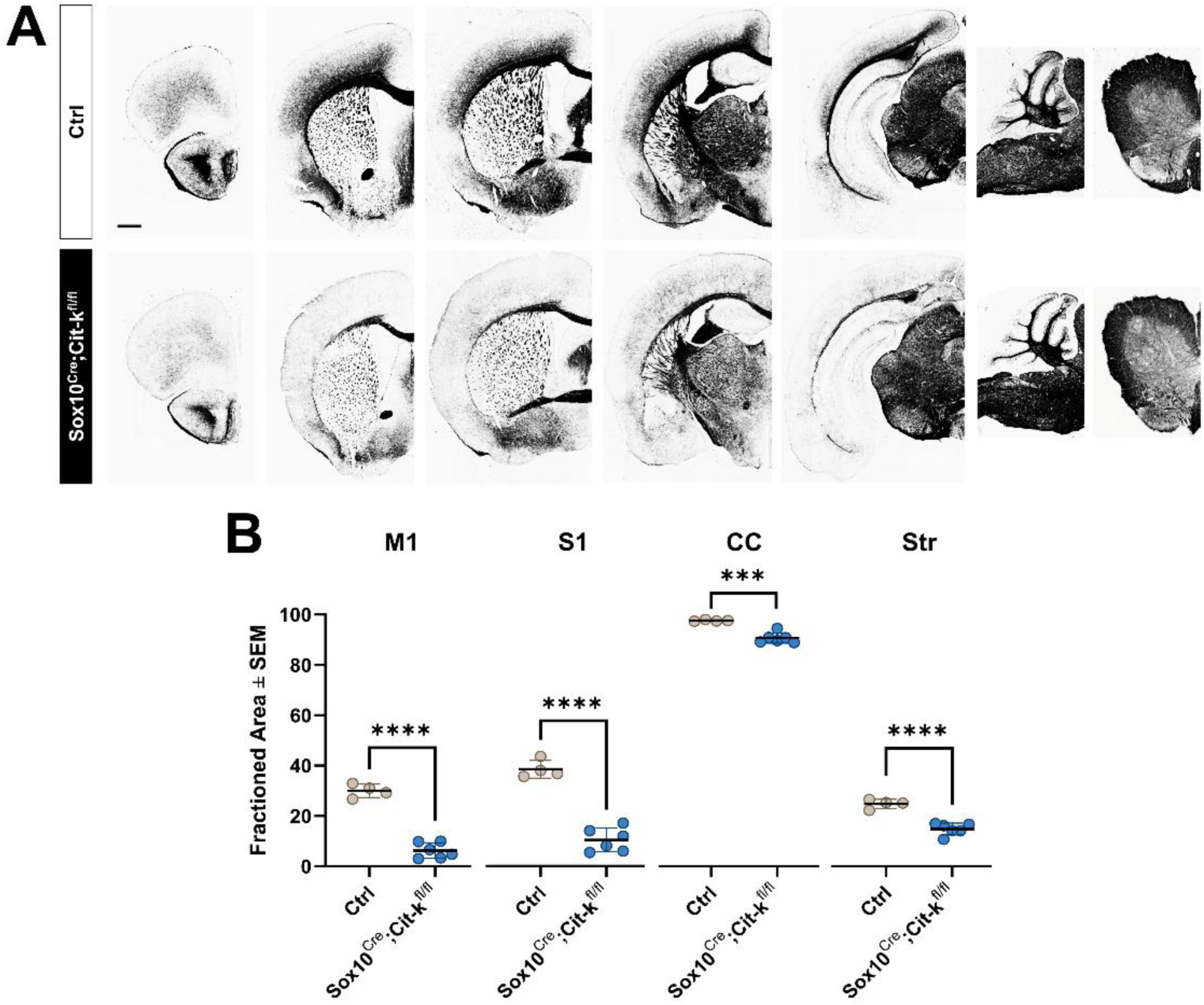
- Gallyas staining confirms persistent cortical hypomyelination in adult Sox10^Cre^;Cit-k^fl/fl^ mice. **(A)** Representative Gallyas myelin staining in coronal forebrain sections at different rostro-caudal levels, sagittal cerebellar and horizontal spinal cord slices (last 2 panels) from adult (4–6 months) control (Ctrl) and Sox10^Cre^;Cit-k^fl/fl^ mice Scale bar: 500 µm for forebrain. **(B)** Quantification of Gallyas-positive area fraction in the primary motor cortex (M1), primary somatosensory cortex (S1), corpus callosum (CC) and striatum (Str). Each dot represents one mouse. Data are presented as mean ± SEM. Statistical significance was assessed by Student’s *t* test (see Suppl.Table 1). ***P < 0.001, **** P<0.0001. Source data are provided in the Source Data file.

**Supplementary Figure 5.**
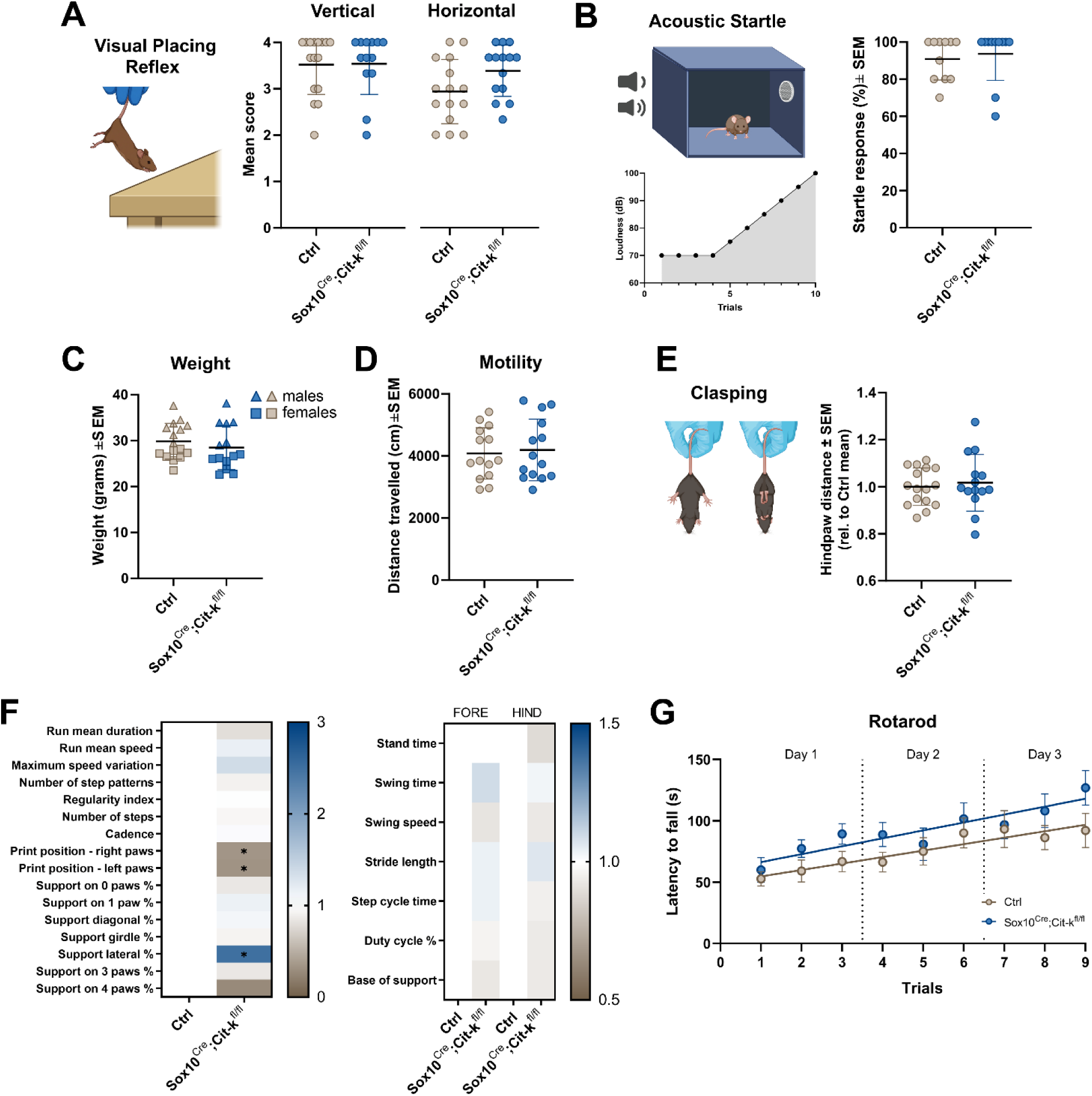
- Preserved sensory function, gross motor performance, and motor learning in adult Sox10^Cre^;Cit-k^fl/fl^ mice. Behavioral assessment of adult (4–6 months) control (Ctrl) and Sox10^Cre^;Cit-kf^l/fl^ mice. **(A)** Visual placing reflex score. **(B)** Acoustic startle response, expressed as the percentage of positive responses to acoustic stimuli of increasing intensity. Graph represents startle responses to 70 dB acoustic stimuli (lowest intensity tested).**(C)** Body weight. **(D)** Spontaneous locomotor activity, expressed as the total distance traveled in an open-field arena. **(E)** Hindlimb clasping behavior, quantified as normalized hindpaw distance during tail suspension. **(F)** Heatmaps of general (left panel) and paw specific (right panel, FORE: forelimb, HIND: hindlimb) locomotion parameters assessed during Catwalk analysis. **(G)** Accelerating rotarod performance across 3 consecutive days (3 trials/day). Latency to fall was averaged across daily trials. Each dot represents one mouse, except in **(F),** where data are presented only as mean ± SEM. Statistical significance was assessed by Student’s *t* test, Mann–Whitney *U* test, or Two-way ANOVA, as appropriate (see Suppl. Table 1). Source data are provided in the Source Data file. Illustrations in A,B,E were created with BioRender.com.

**Supplementary Figure 6.**
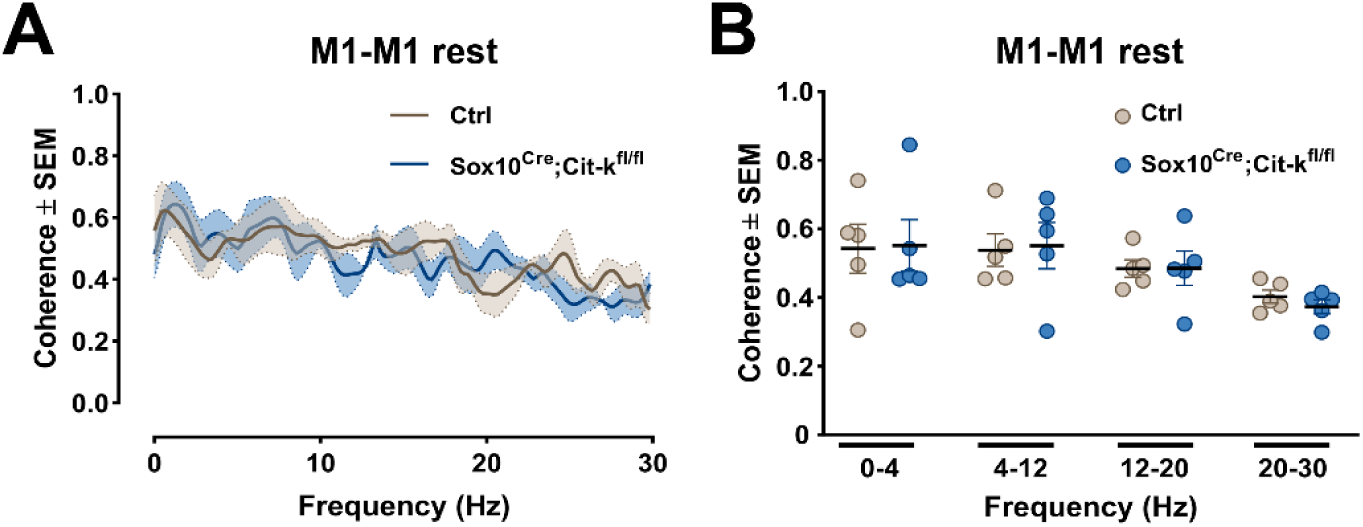
- M1–M1 coherence is preserved at rest in adult Sox10^Cre^;Cit-kf^l/fl^ mice. Assessment of interhemispheric motor cortical synchrony during resting periods. **(A)** Mean coherence spectrum between M1 LFPs recorded simultaneously from the two hemispheres at rest **(B)** Mean M1-M1 coherence values in delta (0.5–4 Hz), theta (4–12 Hz), beta (12–20 Hz), and gamma (20–30 Hz) frequency at rest. Each dot represents one mouse. Data are presented as mean ± SEM. Statistical significance was assessed by Student’s *t* test, Mann–Whitney *U* test, or Two-way ANOVA, as appropriate (see Suppl. Table 1). Source data are provided in the Source Data file.

**Supplementary Figure 7.**
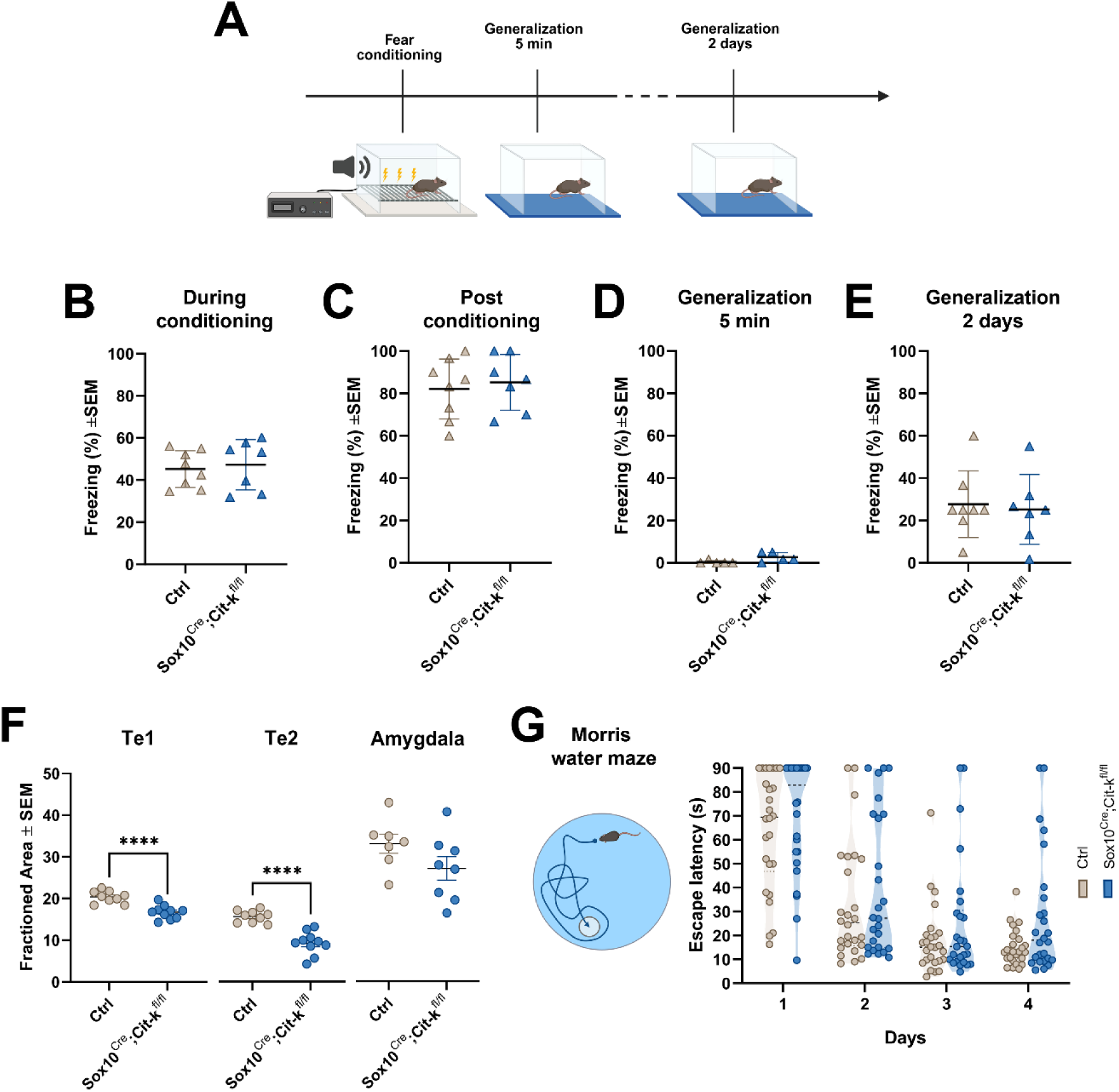
- Preserved fear responses and heterogeneous spatial learning performance in adult Sox10^Cre^;Cit-k^fl/fl^ mice. **(A)** Experimental design of the auditory fear-conditioning protocol used to assess fear acquisition and generalization. **(B)** Percentage of time spent freezing during fear conditioning. **(C)** Percentage of time spent freezing during the immediate post-conditioning period. **(D)** Fear generalization assessed 5 min after conditioning in a novel environment. **(E)** Fear generalization assessed 2 days after conditioning in a novel environment. **(F)** Quantification of MBP-positive area fraction in the primary auditory cortex (Te1), temporal association auditory cortex (Te2) and amygdala. **(G)** Morris water maze performance expressed as escape latency across 4 consecutive training days. Each dot represents one mouse. Data are presented as mean ± SEM, except in (G), where individual data points are overlaid on violin plots illustrating the distribution and variability (i.e. median, quartiles and frequency distribution density of data) of performance within each genotype. Statistical significance was assessed by Student’s *t* test, Mann–Whitney *U* test, or Two-way repeated-measures ANOVA, as appropriate (see Suppl.Table 1). **** P<0.0001. Source data are provided in the Source Data file. Illustrations in A,G were created with BioRender.com.

**Supplementary Figure 8.**
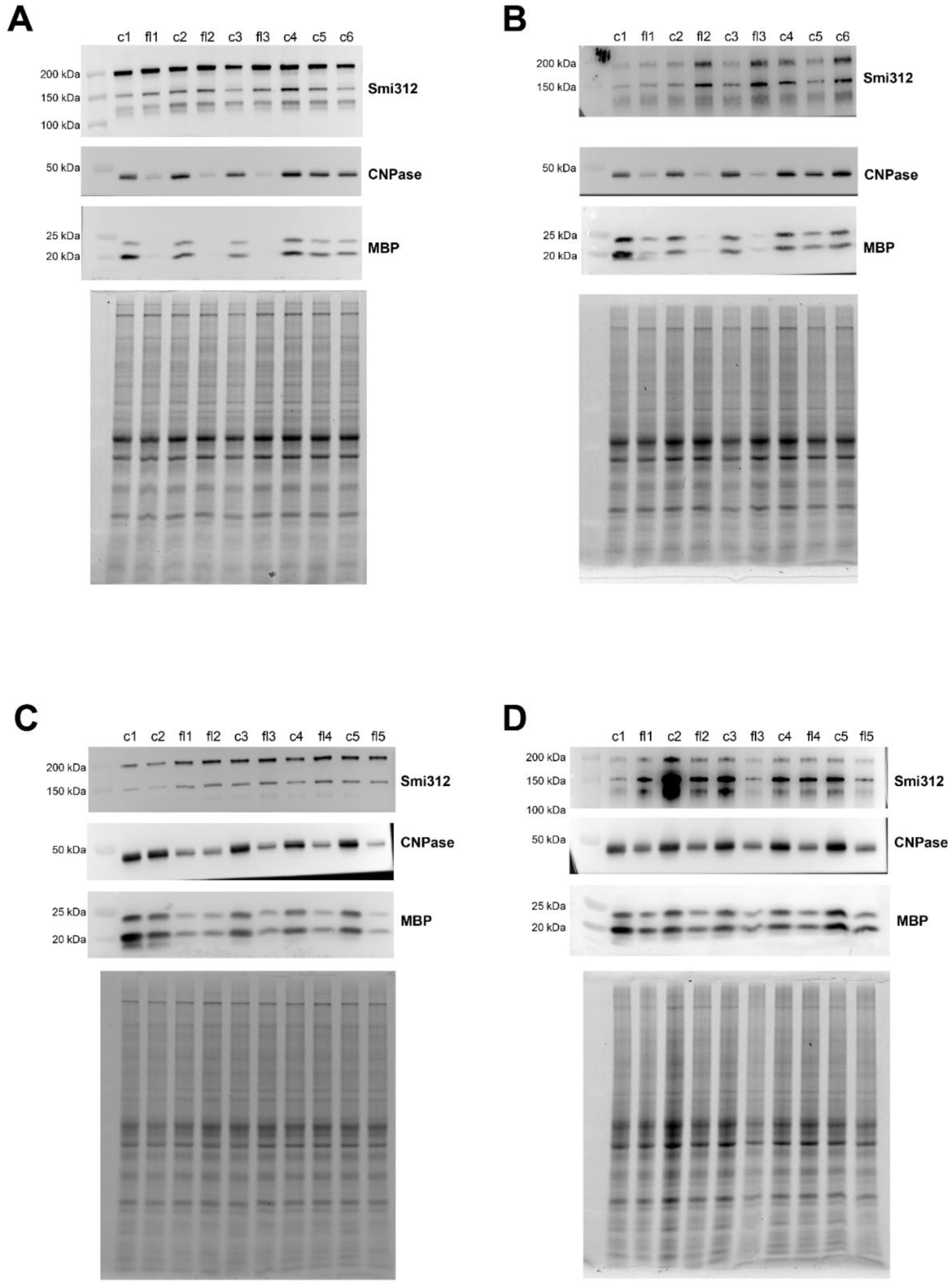
– Original uncropped Western Blot images (from Fig.1C,D and Fig 2D,E) **(A,B)** Western blots of the axonal neurofilament marker Smi312 and of the myelin proteins CNPase and MBP in dorsal **(A)** and ventral **(B)** forebrain lysates from P14 Ctrl and Sox10^Cre^;Cit-k^fl/fl^ mice. **(C,D)** Western blots of the axonal neurofilament marker Smi312 and of the myelin proteins CNPase and MBP in dorsal **(C)** and ventral **(D)** forebrain lysates from adult Ctrl and Sox10^Cre^;Cit-k^fl/fl^ mice. In all cases, TGX stain-free gel imaging (lowest panels) was used as loading control.

**Supplementary Table 1.**
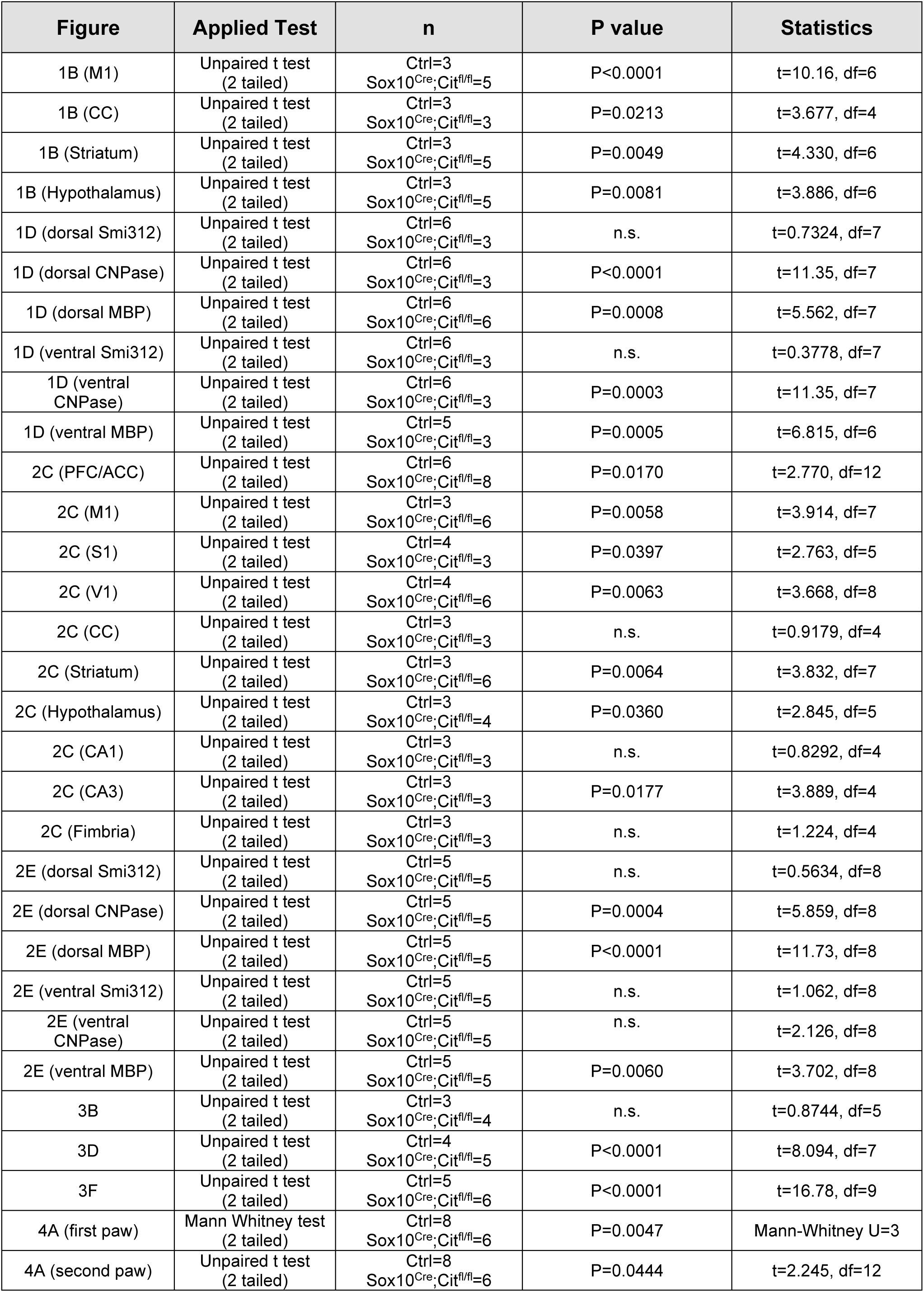
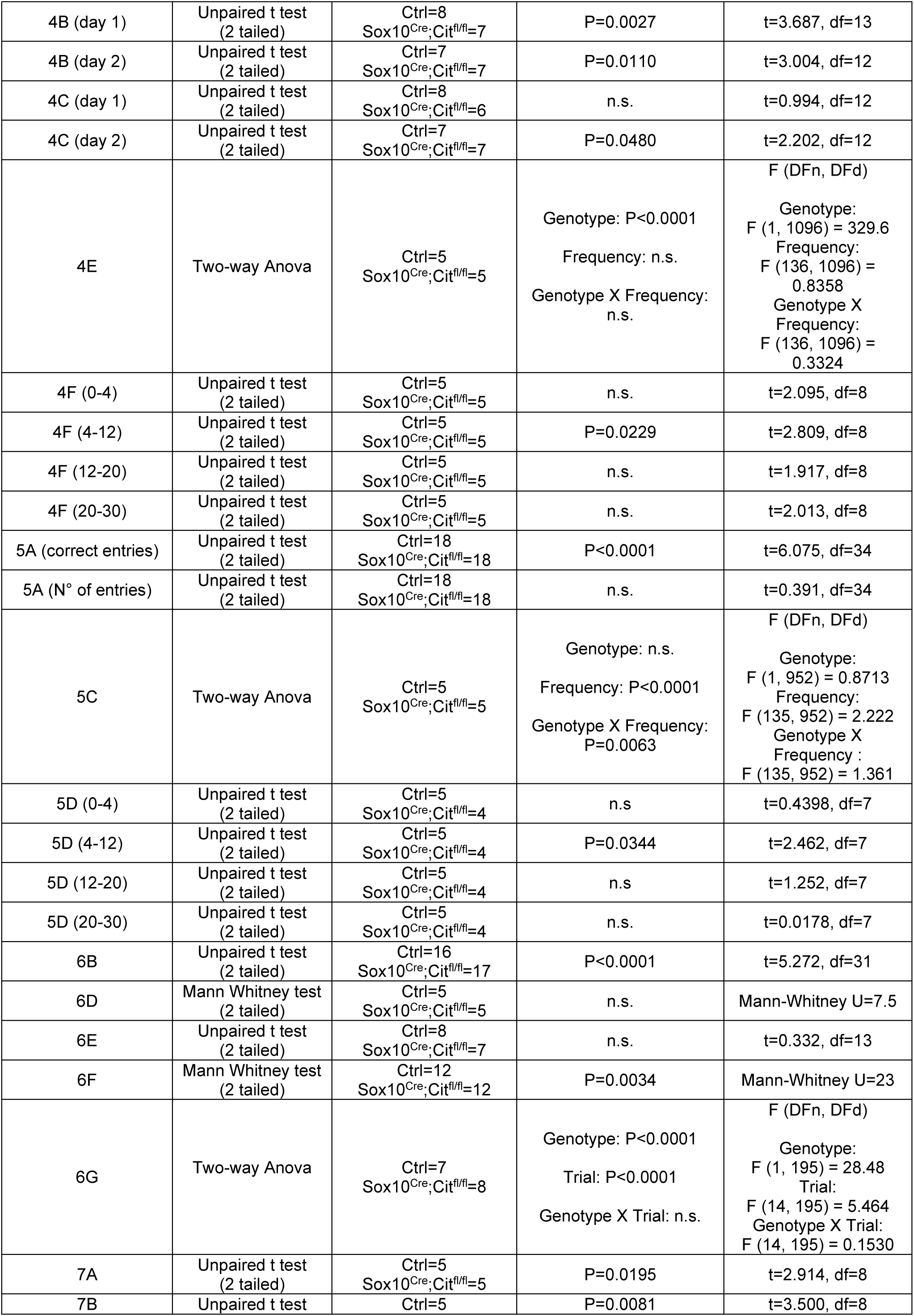
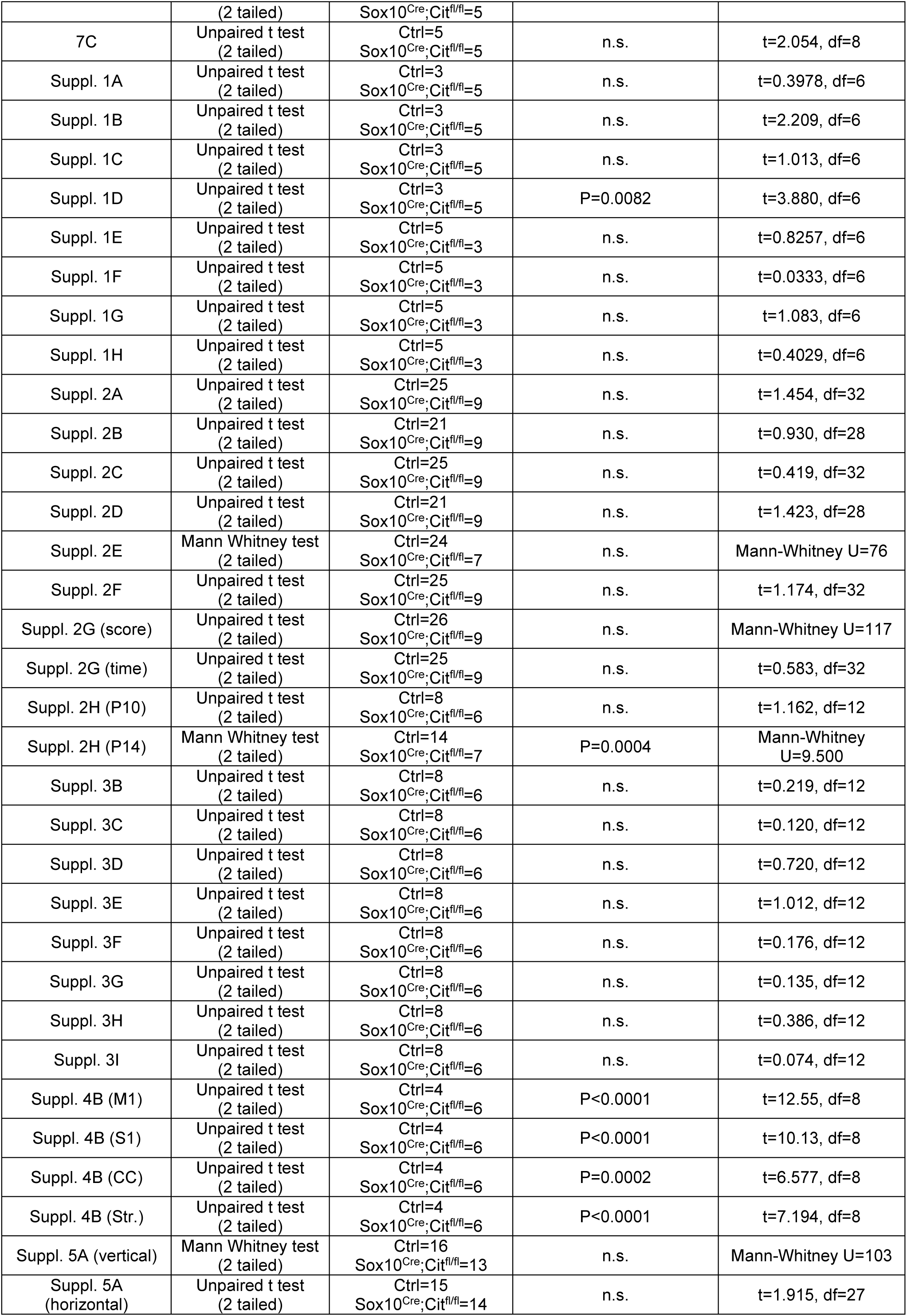
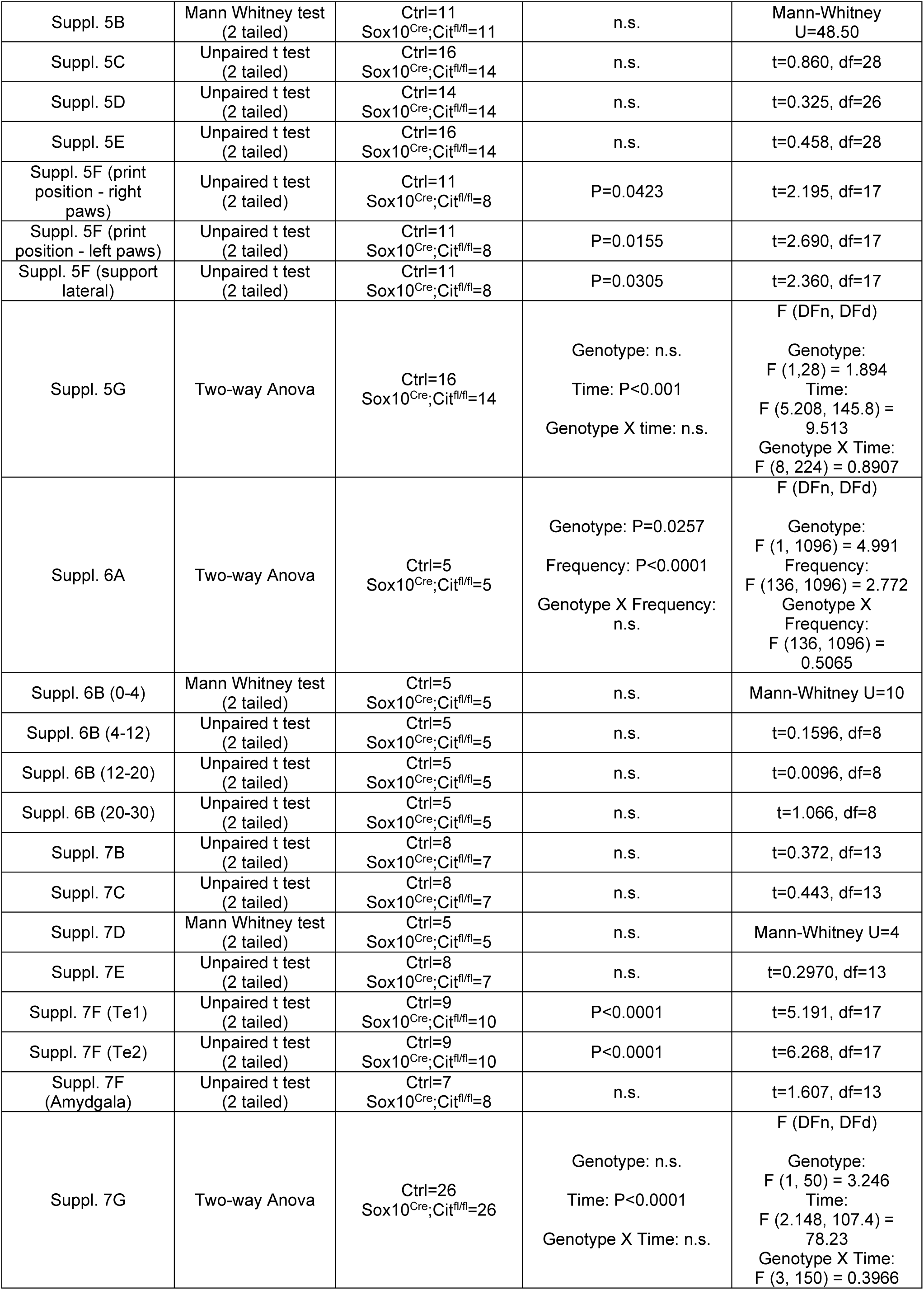
- Statistics.

## Acknowledgements

We wish to thank Dr. Roberta Parolisi and Dr. Giulia Nato (Dept. of Neuroscience Rita Levi Montalcini) for the precious assistance in microscopy analyses. Our work was supported by Telethon Foundation (Multiround 2021-2024 – Round I, ID: GMR22T1066 to EB and Multiround 2025-2027 Round I, ID: GMR25T2067 to EB and MC) and by PRIN (Research Programs of National Interest) 2022 – Italian Ministry of University and Research (ID: 20224YJBBP to EB). This study was also supported by the Italian Ministry of University and Research project “Dipartimenti di Eccellenza 2023–2027” to Department of Neuroscience “Rita Levi Montalcini of the University of Turin. MC was partially supported by the Next Generation EU/Ministry of University and Research project: “A multiscale integrated approach to the study of the nervous system in health and disease (MNESYS)”, CUP B33C22001060002, PE00000006 missione 4, componente 2, investimento 1.3.

## Ethics Statement

This study involves experimental animals and was designed according to the guidelines of the European Communities Council (2010/63/EU) and the Italian Law for Care and Use of Experimental Animals (DL26/2014). It was also approved by the Italian Ministry of Health and conducted according to the ARRIVE guidelines.

## Data Availability Statement

All data are available in the main text and in the Source data file enclosed as Supporting Information.

## Declaration about the use of generative AI and AI-assisted technologies

During the preparation of the manuscript, the authors used ChatGPT to assist with editing for clarity and English grammar after drafting the original outline and manuscript. All scientific content and interpretations were developed independently by the authors, who take full responsibility for the content of the publication.

